# Early migration of precursor neurons initiates cellular and functional regeneration after spinal cord injury in zebrafish

**DOI:** 10.1101/539940

**Authors:** Celia Vandestadt, Gilles C. Vanwalleghem, Hozana Andrade Castillo, Mei Li, Keith Schulze, Mitra Khabooshan, Emily Don, Minna-Liisa Anko, Ethan K. Scott, Jan Kaslin

## Abstract

Zebrafish have a remarkable capacity to regenerate following spinal cord (SC) injury but the responsible cellular events are not well understood. We used *in vivo* imaging and genetics to pin-point specific cellular processes controlling SC regeneration in zebrafish. We identified two temporally and mechanistically distinct phases of cellular regeneration in the SC. The initial phase relies on migration of precursor neurons to the injury, enabling rapid functional recovery, and activation of quiescent neural progenitor cells (NPCs). A second phase of regenerative neurogenesis compensates for both the lost tissue and cells depleted due to precursor neuron migration. We propose a critical role of precursor neurons recruitment in initiating neuronal circuit recovery and buying sufficient time for regenerative neurogenesis to take place. Taken together, our data suggests an unanticipated role of precursor cell recruitment in driving neural repair and functional recovery during the regenerative response.

**Graphical Abstract:** 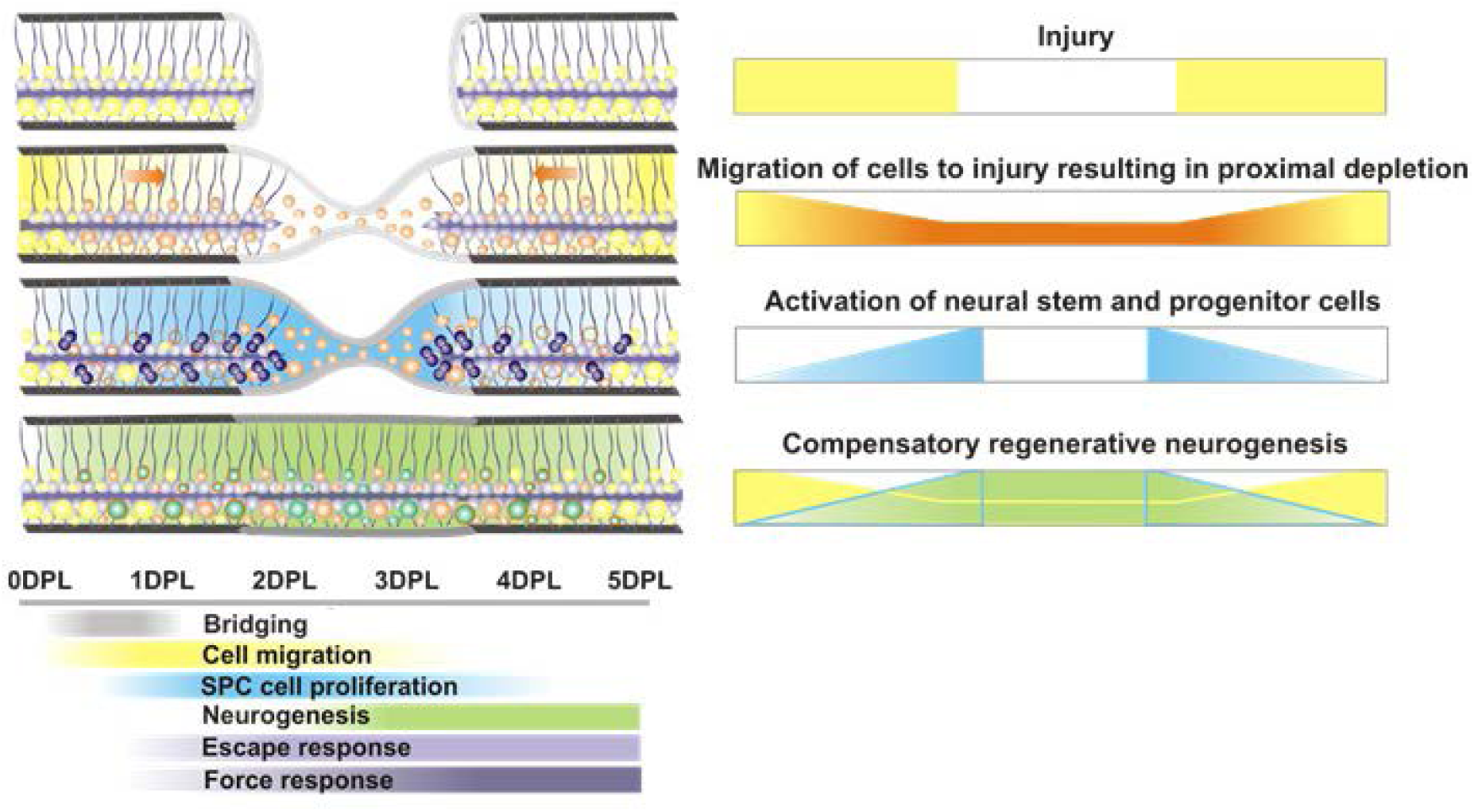

## Introduction

Non mammalian vertebrates, such as teleost fish, can regenerate lost tissue and recover lost function after complete spinal cord injury (Becker et al., 1997, Reimer et al., 2008, Tanaka and Ferretti, 2009, Hui et al., 2010, Kuscha et al., 2012a, Kuscha et al., 2012b, Goldshmit et al., 2012, Briona and Dorsky, 2014). It is well established that many if not all cell types of the spinal cord can be replenished after SCI in zebrafish (Reimer et al., 2008, Hui et al., 2010, Kuscha et al., 2012a, Kuscha et al., 2012b, Briona and Dorsky, 2014). Dormant ependymo-radial glial-like cells (ERGs) around the central canal are activated and proliferate in response to injury, and different ERG domains gives rise to specific subtype of neurons and glia (Grandel et al., 2006, Reimer et al., 2008, Kuscha et al., 2012a, Kuscha et al., 2012b, Goldshmit et al., 2012, Briona and Dorsky, 2014). In particular, motor neuron regeneration has been studied in the larval and adult zebrafish (Park et al., 2007, Reimer et al., 2008, Ohnmacht et al., 2016). Embryonic motor neuron generation from the motor neuron progenitor domain (pMN) largely stops beyond 48 hpf (Reimer et al., 2013), whereas generation of oligodendrocytes from these progenitors continues for many weeks (Park et al., 2007).

Mechanical or genetic ablation injury the adult or larval spinal cord similarly reinitiates the cellular and molecular programmes, including activation of quiescent olig2 expressing ERGs that are required for motoneuron production (Reimer et al., 2008, Ohnmacht et al., 2016). Injury results in regenerative neurogenesis of diverse neuronal lineages including motor neurons in juvenile and adult fish and occurs within days to weeks after injury respectively (Becker et al., 1997, Reimer et al., 2008, Goldshmit et al., 2012, Briona and Dorsky, 2014). However, genetic lineage tracing and cell labelling experiments using thymidine analogues have resulted in a surprisingly low proportion of newly produced neurons at the injury site prior to recovery of motor function (Reimer et al., 2008, Hui et al., 2010, Kuscha et al., 2012a, Kuscha et al., 2012b, Briona and Dorsky, 2014, Briona et al., 2015, Mokalled et al., 2016, Cardozo et al., 2017). This opens up the possibility that the initial tissue recovery may be governed by migration of existing precursor cells that complements, but is distinct from stem cell driven regenerative neurogenesis.

Enigmatic neurons with immature characteristics have been observed in the mammalian (Luzzati et al., 2014, Piumatti et al., 2018, Rotheneichner et al., 2018) and have been proposed to participate in tissue repair in the CNS (Kempermann, 2008, Bonfanti, 2011, Kempermann, 2012). Intriguingly, precursors described as immature neurons in “standby mode” have been reported in the spinal cord of turtle and rat (Russo et al., 2004, Marichal et al., 2009). Additional studies show that cells with similar immature neuronal characteristics are recruited to the site of injury or inflammation in rodents, suggesting involvement in repair (Mothe and Tator, 2005, Danilov et al., 2006, Shechter et al., 2007, Meletis et al., 2008). Recent work in zebrafish shows that neural progenitors directly migrate to the lesion site and participate in repair without undergoing proliferation (Barbosa et al., 2015). These studies highlight the possibility that immature neurons could directly contribute to neural repair in vertebrates. Understanding the role and mechanisms controlling such immature neurons during regeneration is important since these finding could be applied to facilitate spinal cord repair in novel ways.

Here we use larval and adult SC lesion (SCL) models, *in vivo* imaging, behavioural, genetic and pharmacological approaches to investigate spinal cord repair. We find that morphological and functional regeneration occurs rapidly and is tightly correlated, even early after injury. The early regeneration occurs prior to significant neurogenesis and requires existing neurons that border the lesion to migrate into the injury site, resulting in specific compensatory neurogenesis in the uninjured border region. These existing neurons have a distinct identity from both progenitor and mature neurons and we have termed them “precursor neurons”.

## Results

Glial and neuronal cell types regenerate rapidly following spinal cord injury in larval and juvenile zebrafish (Briona and Dorsky, 2014, Ohnmacht et al., 2016). However, the temporal dynamics of different cell types and correlation of functional recovery is not well established. We performed highly reproducible microsurgical spinal cord lesions (SCL) at three days post fertilisation (3dpf, Figure 1A-1C) in transgenic reporter lines to compare the temporal response and regenerative potential of different cell types to injury (Figure 1D-1P). *In vivo* imaging of the regenerative process using a glial reporter *Tg(gfap:GFP*)^*mir2001*^ and pan-neuronal reporter *Tg(HuC:EGFP, HuC* aka *elavl3*)^*knu3*^ showed that both cell types bridge the injured gap within 24 hours after injury demonstrating that glial and neuronal bridging occurs rapidly and simultaneously (white arrowheads, Figure 1E, 1I and Movie S1), consistent with a recent study (Wehner et al. 2017). To examine a distinct population of neurons in the cord we quantified the recovery of the *Tg(isl1:GFP*)^*rw0*^ (referred to as *Islet1*:GFP) cells that labels secondary motoneurons and ventral interneurons (Higashijima et al., 2000). *Islet1*:GFP cells were observed in the lesion site already 1 day post-lesion (DPL, Figure 1M). Recovery of glial and neuronal cells and processes continued rapidly and at 5DPL 96.6%± 0.7%, 83.6%± 2.2% and 59.8%± 5.4% of *gfap*^*+*^, *HuC*^*+*^ and *Islet1*^*+*^ cells, respectively, within the lesion were restored (Figure 1G, 1K, 1O and 1P, for method of quantification see Figure S1).

**Figure 1.**
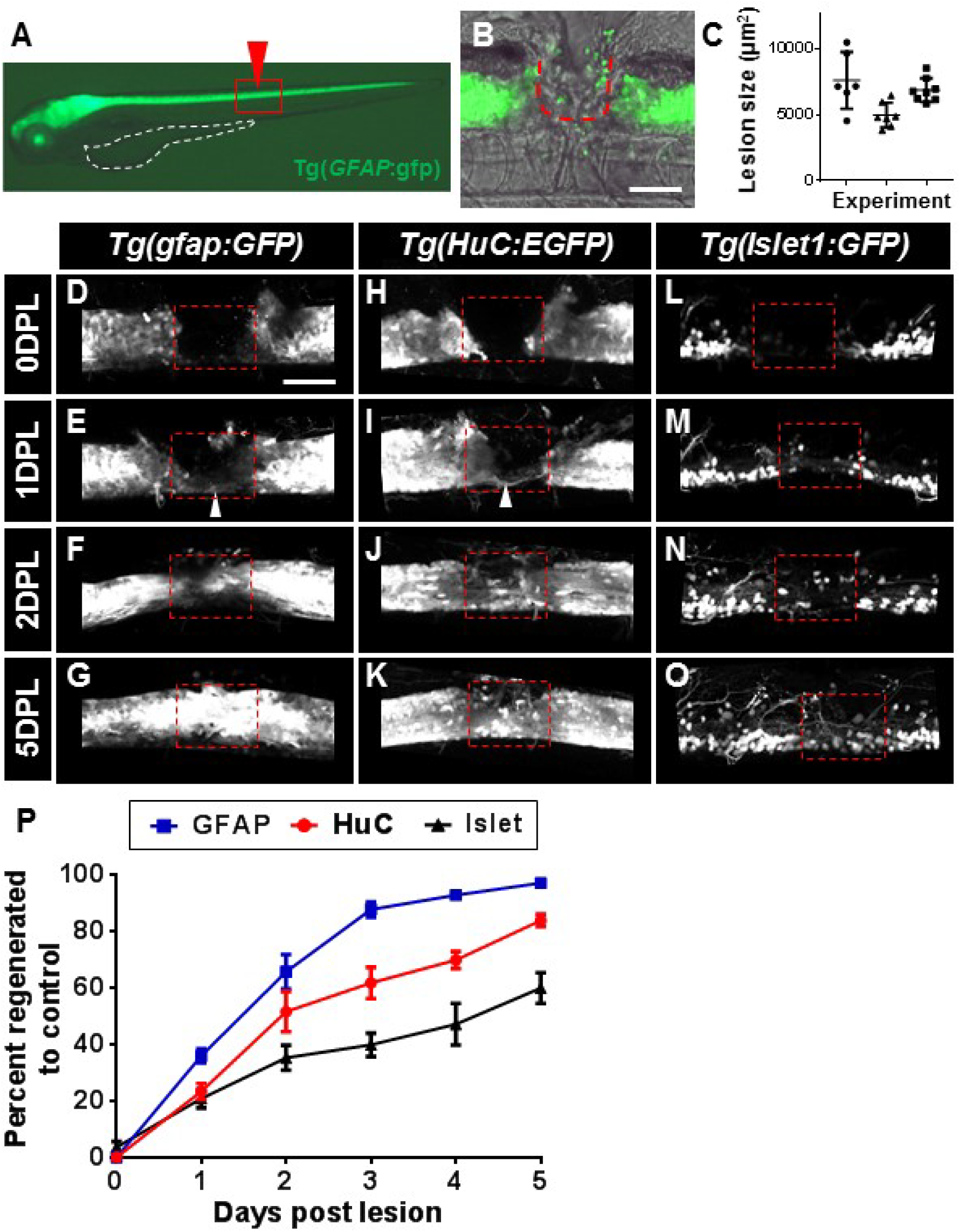
Neuronal and glia cell types have a rapid morphological regenerative response following injury. (A) Lateral view of a *Tg(gfap:GFP*) larva at 3dpf, showing glial cells expressing GFP in the brain and spinal cord, yolk sack outlined in white and red box indicates field of view in B. (B) A precise mechanical lesion at the level of anal pore, indicated by red arrowhead in A, ablates the spinal cord and surrounding tissues, scale bar 100µm. (C) Quantification of lesion area shows high reproducibility of injury size among individuals and experiments. (D) The spinal cord lesion generates a complete transection of glial cell continuity, scale bar 100µm for D-O (E) At 1DPL glial cells start to bridge and connect the spinal cord stumps along the ventral spinal region, indicated by arrowhead. (F) The bridging process continues and at 2DPL 65.5% ±13.6% glial tissue is reconnected. (G) The glial tissue is largely restored at 5DPL. The regeneration process in the pan-neuronal transgenic line *Tg(HuC:EGFP*) shows that after a complete spinal cord transection (H), a thin neuronal bridge forms ventrally within the lesion site at 1DPL (I), the bridge reconnects about 50% of the tissue by 2DPL (J) and the neuronal tissue architecture is significantly regenerated at 5DPL (K). After a complete spinal cord transection (L) a small number of neurons labelled by the *Tg(Islet1*:*GFP*) are visualized within the lesion site (highlighted by red dashed box) at 1DPL (M). At 2DPL an increased number of *Tg(Islet1:GFP*) neurons are located within the lesion site (N) and by 5DPL nearly 60% of *Tg(Islet1:GFP*) cellular regeneration has occurred (O). (P) Quantification of the temporal regenerative response in glial, pan-neuronal and ventral neuron reporter lines. At 5DPL glial cells have completely regenerated, while 83.7% ±6.3% of neurons have recovered and 59.8% ±1.4% of *Tg(Islet1:GFP*) neurons are restored at the lesion site. N=5-11 larvae, error margin = SEM (standard error of the mean), p= ***<0.001, ****<0.0001.

The rapid restoration of lost cell types and processes at the lesion site implied fast recovery of motor functions. To assess locomotor capacity we measured the initial C-bend angle following an elicited startle response (Eaton et al., 1984, Kimmel et al., 1974). Six hours following injury the maximal C-bend angle was reduced by 24%, followed by a return to normal range by 2DPL (Figure 2A and 2B, Movie S2). We next examined the functional engagement of muscles caudal of the lesion site by measuring the active contractile force (Figure 2C). The trunk was externally excited by electrical stimuli with a duration range of 0.1-5ms where 5ms represent the maximal activation of muscle contraction and force generation by the trunk (Figure S2). Lower pulse durations primarily activate locomotor neurons with low excitation threshold and can be used as readout of spinal cord function (Abramsson et al., 2013). At a low to moderate stimuli that mainly activates the motor circuits in the spinal cord the force production was significantly reduced at 6HPL (p-value=0.002), but significantly improved at 2DPL (p-value=0.009), and returned to sham control level at 4DPL (Figure 2C).

**Figure 2.**
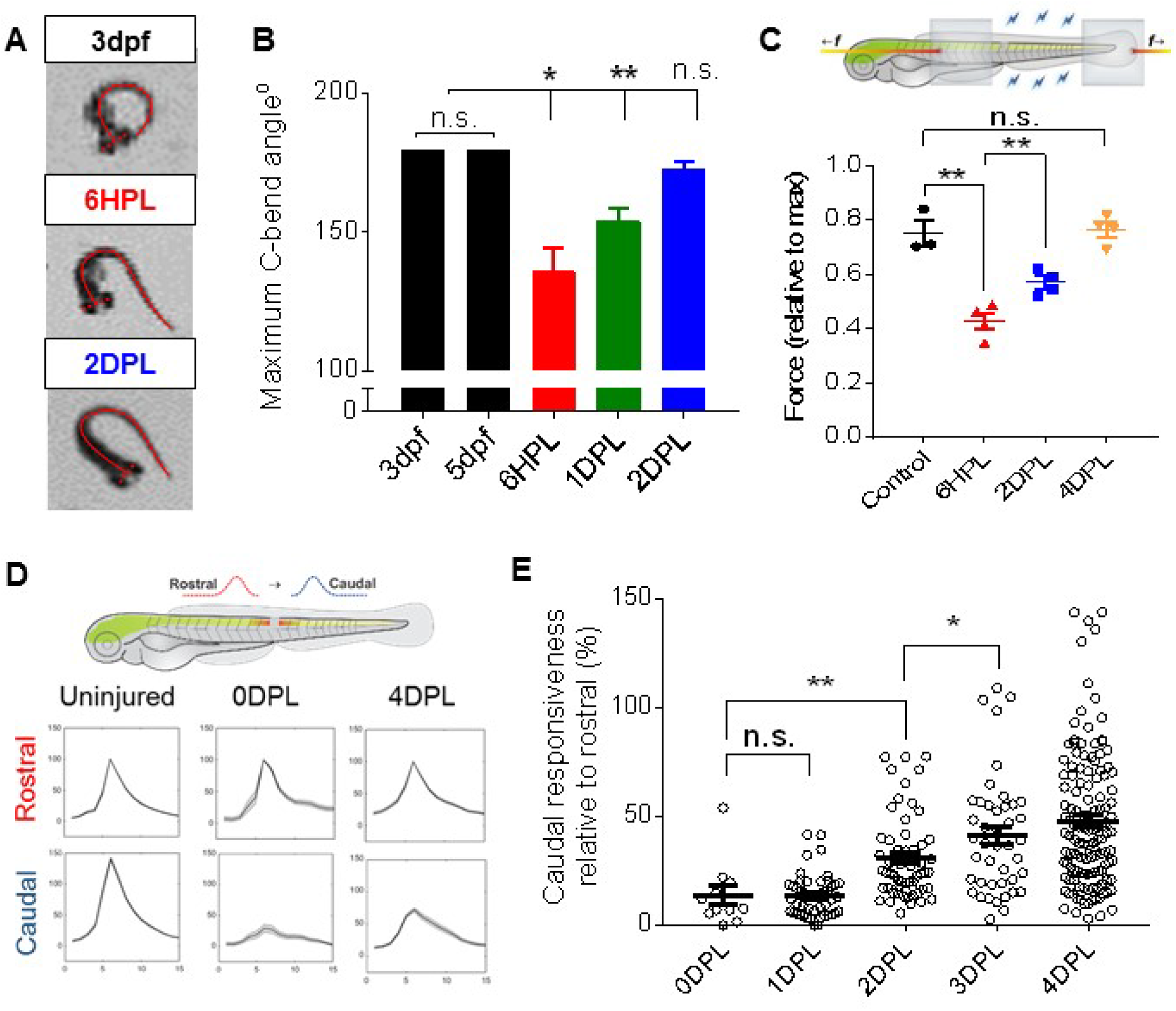
Zebrafish larvae show early functional recovery after spinal cord injury. (A) Dorsal view of the maximal C-bend angle as measured by the angle produced between the head tip to tail tip. Uninjured larvae showed a complete 180° response, while larvae at 6HPL are unable to connect the tip of their tail with their head and at 2DPL locomotor ability is within normal range. (B) Following injury, the maximal C-bend angle was reduced by 24% at 6HPL. At 1DPL the maximal C-bend angle had improved to a 14% reduction compared to unlesioned larvae and was not significantly different to uninjured controls by 2DPL. (C) Schematic of muscle force recording preparation with aluminum clips indicated by grey boxes, between which force measurements are recorded as indicated by blue bolts. Orange lines with f indicate the mechanical pull exerted on larvae to maintain optimal muscle length. At 6HPL force production is significantly reduced compared to uninjured control, while a significant improvement of force production at 2DPL and a non-significant difference between 4DPL and control was observed with a 0.4 ms pulse stimulus. (D) Schematic of GCaMP6s activity propagation analysis. The fluctuation in fluorescence was recorded within both rostral and caudal segments in the 5 seconds before and 10 seconds following a spontaneous motor contraction. Rostral (red) and caudal (blue) mean trace fluctuations are plotted with SEM represented in grey for uninjured, 0DPL and 4DPL. (E) Peak fluorescence scores acquired from method in (D) were used to represent the responsiveness of the caudal segment as a percentage of peak responsiveness in rostral segment for each timepoint following injury. No significant difference in caudal responsiveness was detected at 1DPL compared to 0DPL. Return of % caudal responsiveness significantly increases from 2DPL, but does not reach rostral levels (100%) by 4DPL. N=3-12 larvae, error margin = SEM (standard error of the mean), p= n.s.= not significant, *<0.05, **<0.01. Statistical analysis was measured using students t-test.

To define if functional recovery was detected at the neural circuit level we used diffuse light-sheet microscopy to image neural activity in the spinal cord after injury using GCaMP6s driven by the pan-neuronal promotor HuC/D (Chen et al., 2013, Taylor et al., 2018). All larvae showed spontaneous motor contractions characterised by propagation of Ca2+ pulses along the spinal neurons. Immediately after injury and at 1DPL, Ca2+ pulses associated with motor contractions did not propagate downstream of the injury (Figure 2D). At 2DPL neural activity could be detected in cells at lesion site and Ca2+ waves propagated through the lesion site in a small proportion of larva (data not shown). By 2DPL larvae showed a significant return of GCaMP responsiveness in the caudal stump which continued to improve in the following days, demonstrating rapid recovery of neural circuit activity in agreement with behavioural assays (Figure 2E). Taken together, larva demonstrate a significant recovery of motor function already by 2DPL and in agreement with the rapid recovery of cells and fibres.

The rapid morphological and functional recovery following SCL suggested that regenerative neurogenesis characterised by neural stem cell proliferation and differentiation was activated immediately after injury. To determine the temporal proliferative response of neural stem and progenitor cells after spinal cord injury, we performed short EdU pulse-chase experiments at consecutive days following injury. EdU is a thymidine analogue that is incorporated into DNA during S-phase of cell-cycle and can be administered to specifically label cycling cells. To temporally identify when cells proliferate after injury EdU was administered as a single 2h pulse and larvae were subsequently fixed on each consecutive day following injury (Figure 3A). Initially at 1DPL proliferating cells were distributed more widely throughout the cord structure surrounding the lesion site and at later time-points EdU+ cells were located close to the lesion centre (Figure 3B). Proliferation in the spinal cord was significantly increased from day one to day three after injury and significantly reduced at 4DPL (Figure 3C). These experiments demonstrated that the bulk production of new born cells occurs following 1DPL and takes place during a short temporal window (1-3 DPL) in close proximity to the lesion.

**Figure 3.**
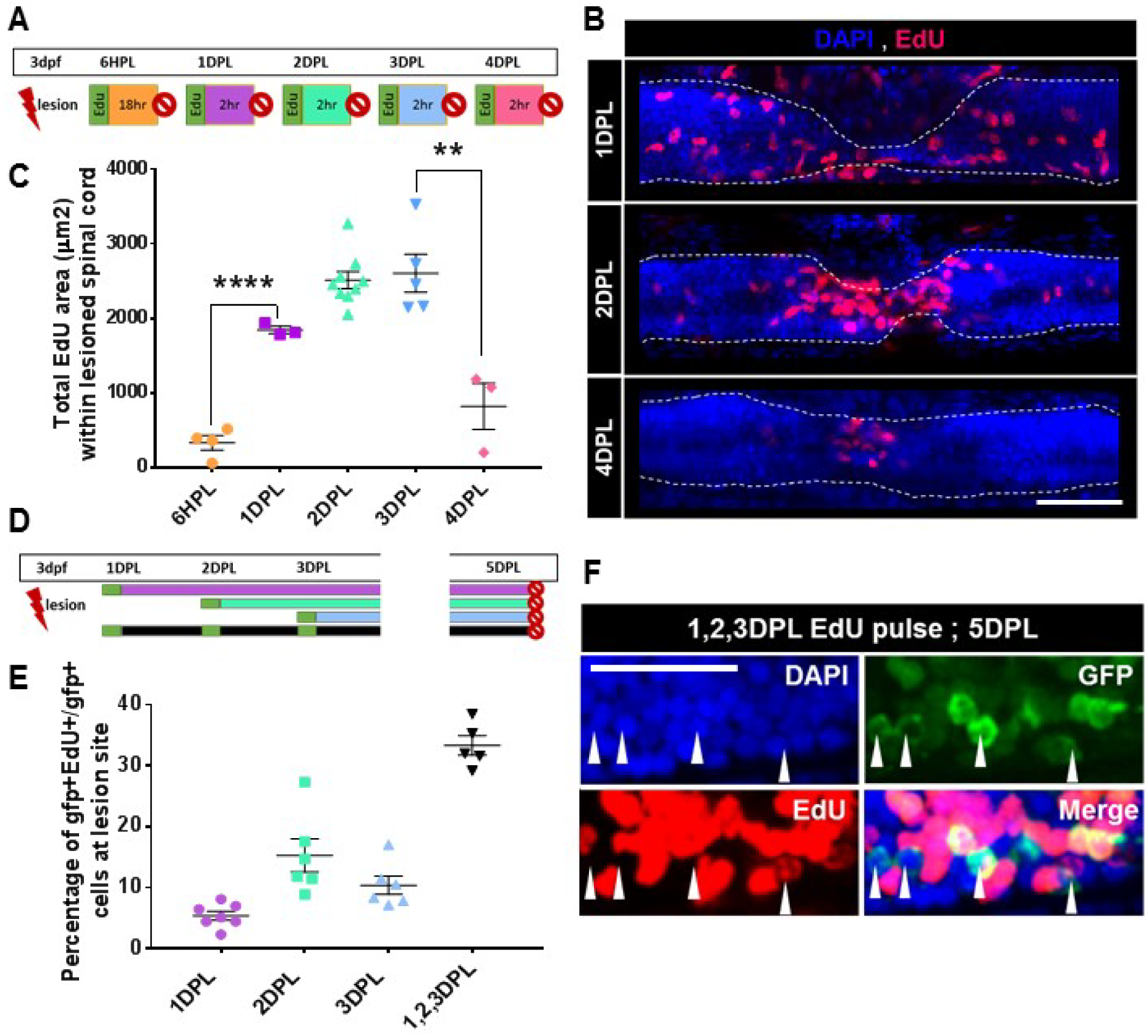
Proliferation and neurogenic response following spinal cord injury. (A) Experimental deign of short pulse/chase proliferation experiment, each day after injury larvae were pulsed with EdU and chased for 2 hours before sacrificing. (B) Representative maximum projections of confocal stacks taken at the lesion site, nuclei are labeled with DAPI and proliferative cells are labeled with EdU (red), showing the dynamics of proliferative cells within the spinal cord. The spinal cord is outlined with white dashed line, as indicated by DAPI nuclei, scale bar 100µm. (C) Quantification of proliferation within the spinal cord at each time point after injury. Proliferation is significantly increased at 1DPL when compared to 6HPL. The peak of proliferation occurs between 2DPL and 3DPL, then proliferation becomes significantly reduced by 4DPL when compared to 3DPL (p-value=0.0047). (D) Experimental design for long pulse/chase neurogenesis experiment in Tg(*Islet:GFP*) larvae, EdU pulses are indicated with green boxes and all larvae were chased until 5DPL. (E) Quantification of *Islet1*:GFP neurogenesis as indicated by the percentage of EdU^+^/*Islet1*:GFP^+^ cells compared to all *Islet1*:GFP^+^ cells within the lesion site at 5DPL. (F) Representative maximal projection of selected confocal stacks from cumulative pulse/chase (indicated by the black bar in D), white arrow heads indicate single positive *Islet1*:GFP^+^ neurons without EdU labelling in the lesion site, scale bar 50µm. n=3-9 larvae, error margin = SEM (standard error of the mean), p= **<0.01, ****<0.0001. Dorsal is up and rostral is to the left in all images.

To temporally identify when neurons were produced after injury larvae were EdU pulsed at specific time-points during the peak of regenerative cell proliferation. To detect when motor neurons and interneurons at the lesion site are produced *Islet1*:GFP larvae were pulsed once with EdU at 1, 2 or 3DPL and fixed at 5DPL for analysis of neurogenesis (Figure 3D). The EdU pulse-chase experiment from 1DPL resulted in very low proportion of *Islet1*:GFP^+^/EdU^+^ cells at the lesion site, only 5.4% (± 1.9%, n=7) of all *Islet1*:GFP^+^ cells co-localised with EdU. Similarly, the Edu pulse-chase experiments from 2 or 3DPL produced a higher, but still a relatively low amount of *Islet1*:GFP^+^/EdU^+^ cells at the lesion site 15.2% (± 6.6%, n=6) and 10.4% (±3.6%, n=6), respectively (Figure 3E). Overall, the proportion of *Islet1*:GFP^+^/EdU^+^ cells compared to single positive *Islet1*:GFP^+^ cells at the 5DPL lesion site was surprisingly low, suggesting that the EdU pulse scheme may be labelling different pools of stem and progenitor cells. To assess if we could label a larger proportion of newly generated *Islet1*:GFP^+^ cells at the lesion site, we performed subsequent cumulative EdU pulses at days 1, 2 and 3DPL and quantified the co-localisation of EdU with *Islet1*:GFP^+^ cells repopulating the lesion site at 5DPL (black bar, Figure 3D). Based on previous studies the cumulative pulses and length is sufficient to capture most if not all cells entering cell cycle and cell cycle time (>14h) is long enough to prevent dilution of EdU label beyond detection limit (Grandel et al., 2006, Kaslin et al., 2009). However, the cumulative pulse chase resulted in only 33% of the *Islet1*:GFP^+^ cells at the lesion site co-localising with EdU (± 3.6, n=5, Figure 3E and F). This was in agreement with the single pulse-chase experiments which quantitatively showed a similar proportion of cells were co-localising (31%) if the proportion of cells from each day were added together (Figure 3E). Taken together, EdU pulse chase experiments after injury showed that while a substantial amount cells are produced, neuronal differentiation of de novo produced cells takes time and mechanisms other than neurogenesis contribute to cellular recovery at the lesion site.

To determine if similar mechanisms of cellular recovery operate in the adult, we performed spinal cord injury in adult *Islet1*:GFP fish and performed consecutive cumulative EdU pulses at 3, 5 and 7 DPL and chased until 18 DPL (Figure 4A). In agreement with the larval data, many of the *Islet1*:GFP^+^ neurons within the lesion centre were not labelled with EdU after injury (clear arrowheads, Figure 4C-D). This is in agreement with previous studies in adult zebrafish showing a substantial increase (>30 fold) of *Islet1*:GFP^+^ or *Hb9*:GFP^+^ expressing cells at lesion site 1-2 weeks after injury that lacked BrdU labelling despite receiving multiple pulses after injury (Reimer et al., 2008). Taken together, a low amount of neurons at the lesion site were co-labelled with EdU even after multiple EdU pulses during the temporal window when most cells proliferated in the spinal cord suggesting that alternative mechanisms, in addition to cell proliferation driven neurogenesis, play a role in the initial cellular and functional recovery. Consistent results within larvae and adult suggest that mechanisms of cellular regeneration are conserved across the lifespan.

**Figure 4.**
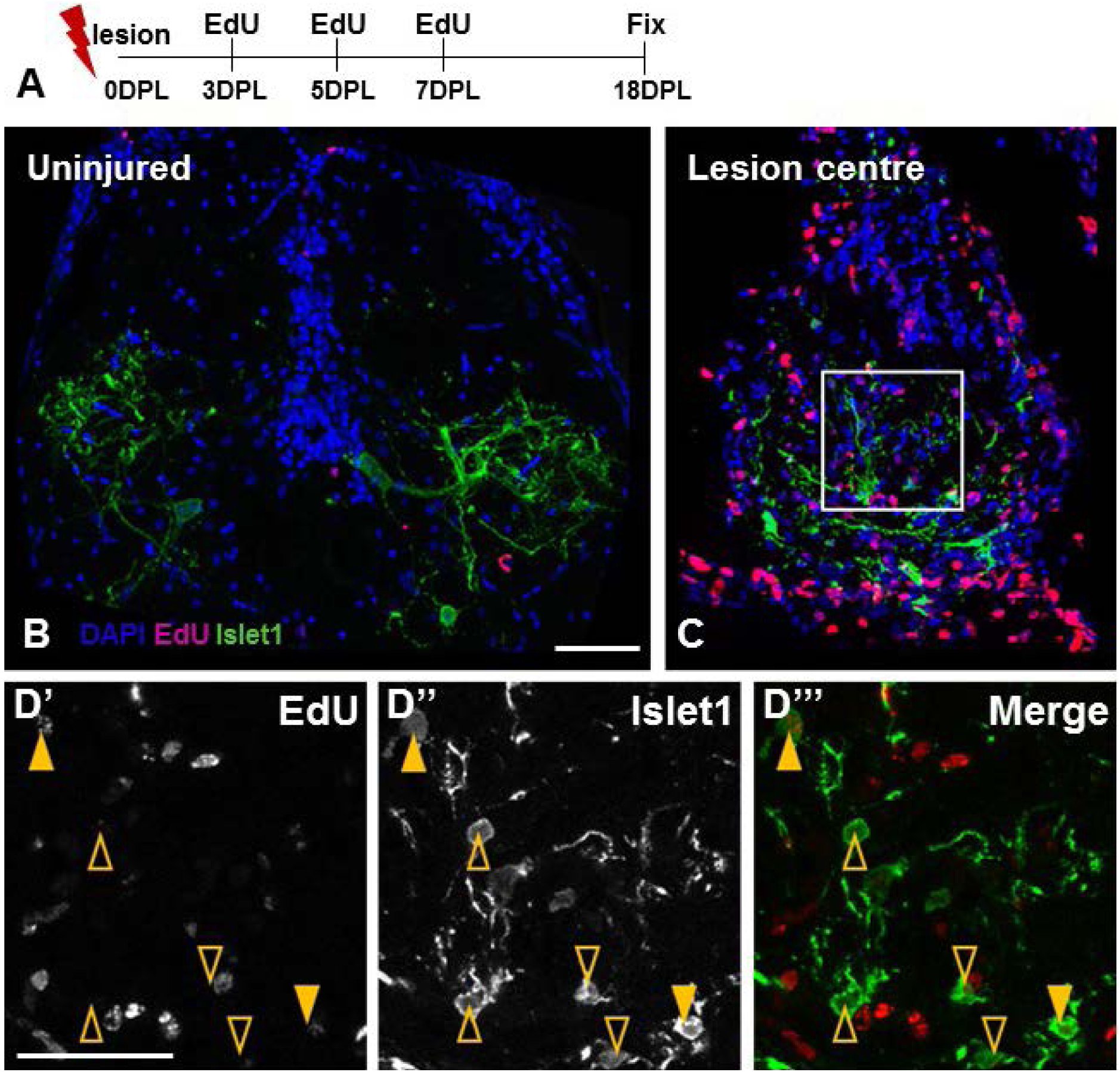
Early neurogenic response following adult spinal cord injury. (A) Schematic of experimental design for adult SCL neurogenesis assay. Injured and sham control fish were pulsed with EdU at 3, 5 and 7DPL via IP injection, then fixed at 18DPL and processed for EdU staining. (B) Cross-section of a sham *Tg(Islet1:GFP*) adult spinal cord pulsed with EdU. Minimal proliferative cell are observed within the uninjured spinal cord. (C) Cross-section of the injured spinal cord at 18DPL shows an upregulation of proliferative cells within the lesion centre. White box indicates region of high magnification in (D). (D) Many *Islet1*:GFP^+^ cells are observed, but few co-localise with the proliferative marker. Solid yellow arrowhead indicates *Islet1*:GFP^+^/EdU^+^ cells, while open arrowhead indicates *Islet1*:GFP single positive cells. All images are maximum projection of confocal stacks, scale bars 50µm.

Substantial morphological and functional recovery occurs prior to peak cell proliferation and neurogenesis suggesting that other tissue remodelling mechanisms may be activated initially following injury. To determine requirement of proliferation for early cellular and functional recovery, we treated injured fish with the cyclin dependent kinase inhibitors Olomoucin and Ryuvidine to block cell cycle entry during the time-window when neural progenitors start proliferating after injury (1-2 DPL, Figure 5A). Treatment with the cyclin dependent kinase inhibitors efficiently blocked cell proliferation following tail injury (Figure S3A and B). Following treatment after SCI the number of *Islet1*:GFP^+^ cells within the lesion site at 2DPL was unchanged, showing that the initial cellular regeneration of neurons at the lesion site is independent of cell proliferation (Figure 5A-C).

**Figure 5.**
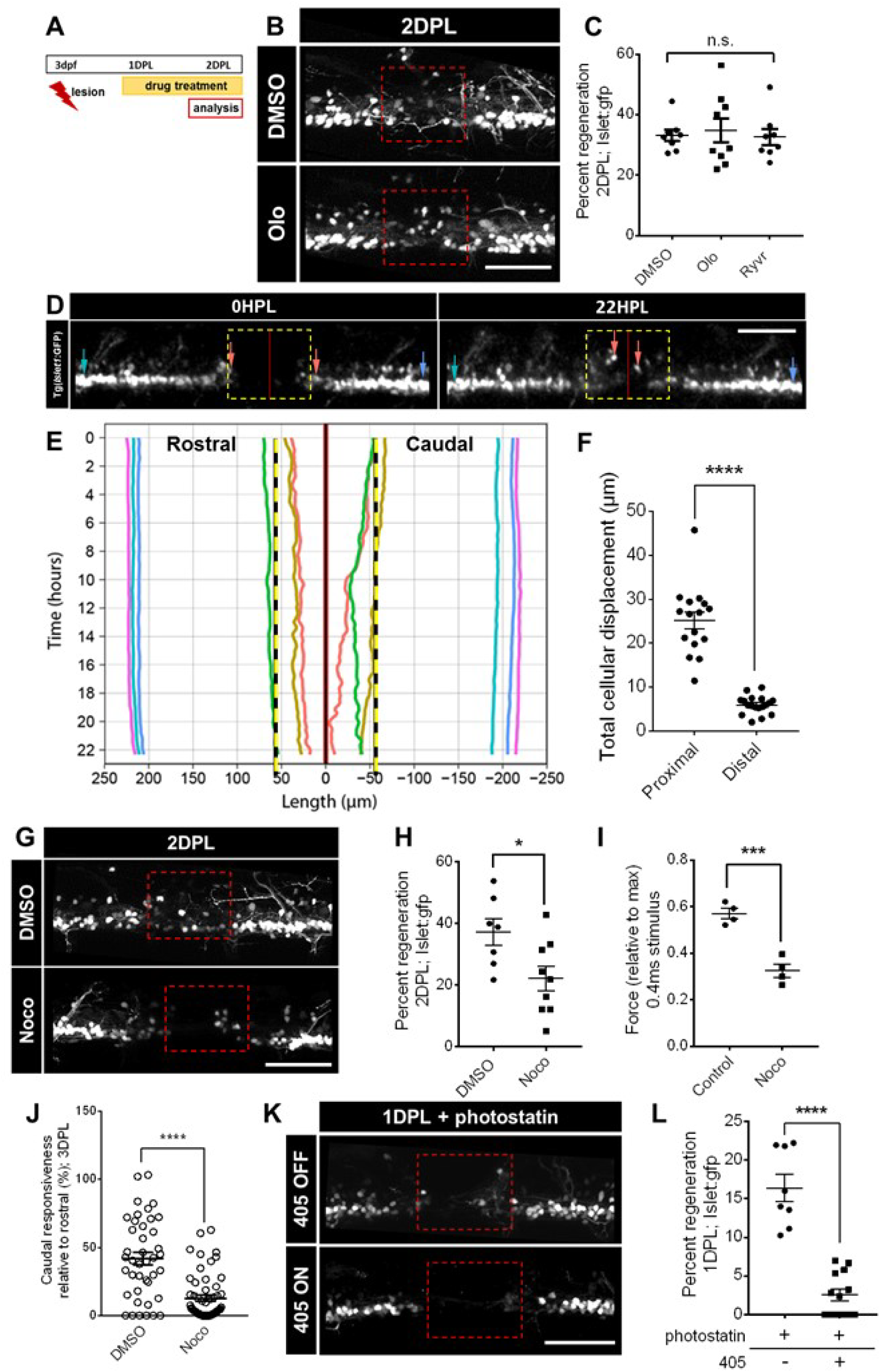
Early neuronal and functional regeneration requires migration, but not proliferation. (A) Scheme depicting experimental design for measuring *Islet1*:GFP regeneration following proliferation inhibition. *Islet1*:GFP larvae were lesioned at 3dpf, drug treatment occurred between 1DPL to 2DPL and larvae were imaged at 2DPL. (B) Maximum projection images of control and Olomoucine treated *Tg(Islet1:GFP*) larvae showing similar number of *Islet1*:GFP^+^ cells in the lesion site. (C) Treatment with cyclin dependent kinase inhibitors, Olomoucine and Ryuvidine, did not impact *Islet1*:GFP^+^ cellular regeneration at 2DPL. (D) *In vivo* maximum projections of confocal stack of the cells surrounding the lesion site in a *Tg(Islet:GFP*) larvae at 0HPL and 22HPL. Individual cells traced in (E) are indicated by corresponding colored arrows. (E) Individual tracks of selected cells from timelapse in (D). Time in hours is displayed on the Y axis and distance in µm along the length axis of the spinal cord relative to the lesion center is shown on the X axis, positive values indicate rostral and negative values indicates caudal with 0 indicating the lesion center. (F) Cells within the proximal segment had a significant increase in overall displacement between 0 and 22HPL compared to cells in distal segments, n=3-4 cells per segment from individual 4 larvae. (G) Maximum projection images of control and Nocodozole treated *Tg(Islet1:GFP*) larvae at 2DPL. (H) Nocodozole treatment, following schematic in (A), reduced the number of *Islet1*:GFP^+^ cells in the lesion site compared to control at 2DPL. (I) Using a stimulus of 0.4ms a 43% reduction in relative force production was recorded in larvae treated with Nocodozole compared control at 2DPL. (J) GCaMP6s peak caudal responsiveness was significantly reduced in larvae treated with Nocodozole compared to control at 3DPL. (K) Representative maximal projection of Tg(*Islet1:*GFP) larvae at 1DPL treated with photostatin, with and without 405 laser excitation, showing an absence of cells in the lesion site in photostatin treated larvae with simultaneous 405 laser excitation. (L) Percent regeneration of *Islet1*:GFP^+^ cells at 1DPL is significantly reduced in larvae treated with photostatin and simultaneously targeted with 405 laser excitation compared to larvae treated with phototstatin alone. N=4-12 larvae, error margin = SEM (standard error of the mean), p= *<0.05, ***<0.001, ****<0.0001. Red boxes indicate lesion site, dorsal up and rostral left in all images, scale bars 100µm.

We next hypothesised that cell migration to the lesion could play an important role in early tissue and functional recovery, prior to the establishment of new born neurons at the lesion site. To capture the cell behaviour and migration of cells after SCI, we used confocal time-lapse microscopy. Imaging of *Islet1*:GFP fish after injury showed that cells in proximity to the lesion boundary migrated towards the lesion centre (orange arrows, Figure 5D and Movie S3). To define individual cell movements we developed an algorithm that measures cell displacement along the length axis of the spinal cord in relation to the lesion centre (Figure S4). Single cells within proximal and distal segments were selected and their trajectories were tracked over time (Figure 5E). The tracking of single cells demonstrated that *Islet1*:GFP^+^ cells proximal to the lesion moved significantly more towards the lesion centre than *Islet1*:GFP^+^ cells that where distally located (p-value<0.0001, Figure 5F).

To determine if cell migration was required for early *Islet1*:GFP^+^ cellular regeneration, we treated injured larvae with Nocodazole, a drug that de-stabilises the polymerisation of microtubules and results in altered cell migration when used at low concentrations (Liao et al., 1995, Baudoin et al., 2008). Treatment of larvae with low dose of Nocodazole after SCI during short time window (between 1 and 2DPL, Figure 5A) significantly reduced the number of *Islet1*:GFP^+^ cells at the lesion site at 2DPL (p-value=0.02, Figure 5G and H). To determine if early *Islet1*:GFP^+^ cell migration impaired functional recovery, we treated larvae with Nocodozole and measured the recovery of contractile force. Treatment with Nocodazole showed no functional recovery at 2DPL compared to vehicle treated larvae (Figure 4I, p-value=0.0005 and Figure S3C). In agreement, GCaMP6s imaging after Nocodazole treatment showed that treated fish displayed a reduced Ca2+ wave response downstream of the injury consistent with observed reduction of recruited neurons (Figure 5J). Systemic treatment with Nocodazole may result in unpredictable general off target effects. To achieve temporal and spatial control of cell migration we used the photoactivatable microtubule inhibitor photostatin (Borowiak et al., 2015). Larvae were treated with photostatin after injury and a restricted volume surrounding the lesion site was subjected to photoactivation at the lesion stumps using low intensity laser pulses at 405nm. Fish that were subjected to photoactivation after SCI had significantly reduced *Islet1*:GFP^+^ cell number at the lesion site at 1DPL compared to larvae treated with photostatin alone (p-value<0.0001, Figure 5K and L).

We next identified the nature and characteristics of cells capable of early migration following injury. As the spinal cord is comprised of distinct dorsal and ventral domains, we first determined if ventral and dorsal spinal cord cell lineages are equally recruited to the lesion. To image ventral cell lineages we used Tg(*olig2*:EGFP)^zf532^ and Tg(*-3.0mnx1*:mtagBFP, mnx1 aka Hb9)^mq10^ that label ventral neural progenitors and motor neurons, respectively (Don et al., 2017). To determine the recruitment of a dorsal population of neurons we labelled 2DPL larvae with pax2. Live imaging of *olig2*:EGFP and *mnx1*:BFP transgenic lines at 1DPL showed that both cell populations rapidly migrated to the injury site (Figure 6A). This was further supported by immunocytochemistry showing mnx1(Hb9)^+^ and Islet1/2^+^ neurons within the lesion site (Figure S5A). Imaging of pax2 stained larvae showed no recruitment of dorsal cell lineages to the lesion site 2 DPL (Figure 6B). These experiments show that ventral neuronal subpopulations preferentially migrate to the lesion early after injury.

**Figure 6.**
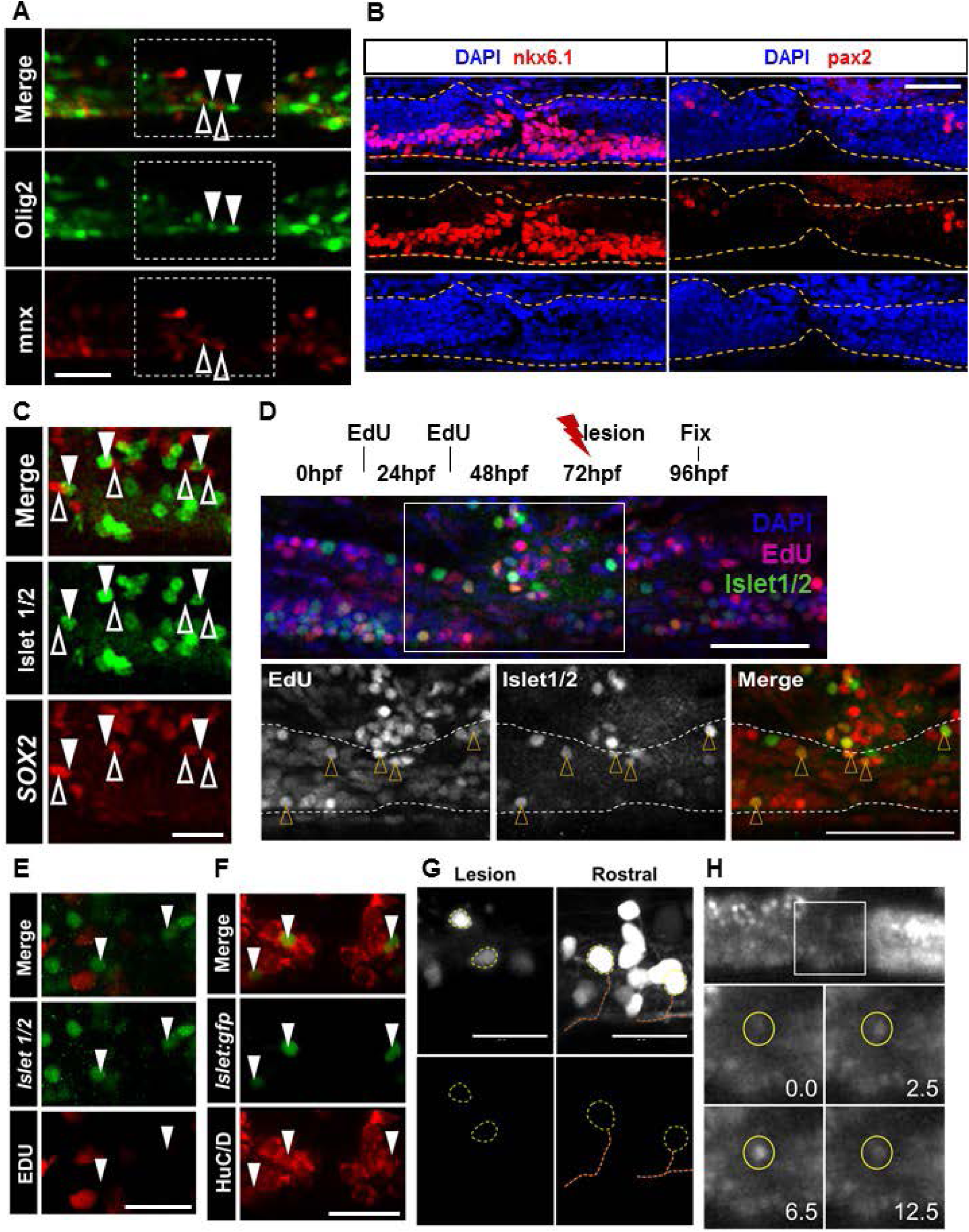
Characteristics of early recruited neuronal precursor cells are distinct from both NSP cells and mature neurons. (A) Maximum projection 1DPL larvae expressing both *olig2*:EGFP and *mnx1*:BFP transgenes. Merge panel shows cells within the lesion site express a singular transgene. Downward facing arrowheads indicate *olig2*:EGFP^+^ cells, while upward facing open arrowheads indicate *mnx1*:BFP^+^ neurons. (B) Maximum projection 2DPL immunostained larvae labelled with either ventral marker nkx6.1 or dorsal marker pax2 showing expression of nkx6.1+ cells but not pax2+ cells within the lesion centre. Spinal cord dorsal and ventral boundaries outlined by orange dotted line, determined by DAPI expression. (C) Maximum projection of 1DPL larvae expressing the *Islet1*:GFP^+^ transgene co-stained for Sox2 expression. *Islet1*:GFP^+^ cells within the lesion site (downward facing solid arrowheads) do not co-localise with the SOX2 marker (upward facing open arrowheads). (D) Experimental design for labelling developmentally derived neurons. Embryos were continuously pulsed with EdU at 14 somite and 32hpf phenotypic stages prior to injury at 72hpf, then fix at 96hfp (1DPL) and processed for staining. Maximum projection overview of lesioned spinal cord at 1DPL in DAPI, EdU and Islet1/2 stained larvae, white box indicates lesion site. Higher magnification of lesion site with spinal cord outlined by white dashed line. Most Islet1/2 positive cells within the lesion centre co-localise with EdU, indicating they were developmentally derived cells, *Islet1*:GFP^+^/EdU^+^ cells are indicated by open orange arrowheads. Scale bar = 100µm. (E) Selective plane maximum projection of 1DPL larvae following short 2hour pulse with EdU and stained for *Islet1/2*. *Islet1/2*^*+*^ cells within the lesion site do not co-localise with the proliferative marker. (F) Maximum projection of 1DPL larvae expressing the transgene and stained for HuC/D, all cells co-localise with the marker for early neuronal fate. (G) In vivo maximum projection of *Islet1*:GFP^+^ cells within the lesion site and rostral spinal segment. Lesion site cells appear smaller than cells located within intact rostral segments, as indicated by yellow dotted lines. Lesion site cells lack processes which is observed by rostral cells, as indicated by orange dotted line. (H) Maximum projection of *HuC/D*:GCaMP6s larvae at 2DPL, response to spontaneous motor contraction. Top panel shows spinal cord one frame after contraction with lesion centre outlined in white box indicating region of magnification for bottom panels. Bottom panels are frames following the contraction with the responsive cell outlined by yellow circle, peaking in fluorescence 6.5 seconds after initial motor contraction. Dorsal up and rostral to left in all panels. Scale bars A, B and D = 100µm, C, E-G = 20µm.

We next determined if early recruited cells showed signs of de-differentiation by expression of neural stem/progenitor or cell proliferation markers. Time-lapse analysis of the *HuC* and *Islet1* reporter lines after injury suggested that cells maintain their specific lineages, since we did not detect cell divisions or appearance/disappearance of reporter expressing cells at the lesion site throughout the time-lapse series (Movie S3). Furthermore, we did not detect expression of the neural stem/progenitor marker Sox2 by recruited *Islet1*:GFP^+^ cells at the injury site 1DPL (Figure 6C). To determine if recruited cells were of developmental origin, we consecutively pulsed embryos with EdU at 16 and 32hpf prior to lesioning to label embryonically derived cells. Immunostaining at 1DPL showed the majority of Islet1/2^+^ cells in the lesion site co-localised with EdU cells that were produced in the early embryo (open arrowheads, Figure 6D). Suggesting that most of the Islet1/2^+^ cells recruited to the lesion site early after injury were existing prior to injury and produced during embryonic development. In agreement, Islet1/2^+^ cells recruited to the lesion site at 1DPL were shown to be non-proliferative, as these cells did not co-localise with the EdU proliferative marker when a short pulse was administered following injury (Figure 6E). However, the recruited Islet1:GFP cells expressed the early neuronal marker HuC/D at 1DPL identifying them as neurons (Figure 6F). Closer examination of the recruited neurons using the *Islet1*:GFP^+^ line showed that cells were small sized, ovoid and lacked elaborate long dendrites or axons, consistent with the morphology of immature neurons (Figure 6G). These experiments showed that the early recruited cells were produced prior to injury and are post-mitotic, but immature within the ventral neuronal lineage.

We next hypothesised that the recruited neurons may act as early pioneers of the regenerating spinal circuitry. Imaging of GCaMP6s in recruited neurons at lesion site showed that the cells started to display coordinated, but temporally delayed neural activity in response to spontaneous motor contractions at 2DPL (Figure 6H). These experiments demonstrated that precursor neurons recruited to the lesion site are the earliest functional pioneers within the lesion site after injury.

A hallmark in regeneration is the ability of injured tissue to identify a loss of cells. Therefore, we next addressed if the cellular depletion due to migration at the proximal regions resulted in secondary injury that was sensed by the quiescent NSCs in the ependyma. We hypothesised that the range of neural stem and progenitor cell activation may correspond to the range of cell depletion which occurs during the early regenerative phase. To determine if existing neurons within the spinal cord regions proximal to the lesion become depleted after injury, we quantified the number of *Islet1*:GFP^+^ cells within increasing distances from the lesion site at 2DPL, a time point when many *Islet1*:GFP^+^ cells were detected at the injury site and prior to significant neurogenesis (yellow boxed area, Figure 7A). Quantification of *Islet1*:GFP^+^ cells demonstrated that the number of cells in the proximal segments was significantly decreased compared to more distal segments (Figure 7B, p-value<0.0001 rostral, p-value=0.0006 caudal). Furthermore, when the number of cells in the lesion site were added together with the cells from proximal segments the cell count was similar to distal unperturbed segments (p=not significant, Figure 7C), suggesting that *Islet1*:GFP^+^ cells were recruited to the lesion from the proximal stumps, resulting in a local cellular depletion. The range of cell depletion was in agreement the recruitment range identified by the live imaging and cell tracking experiments (Figure 5E).

**Figure 7.**
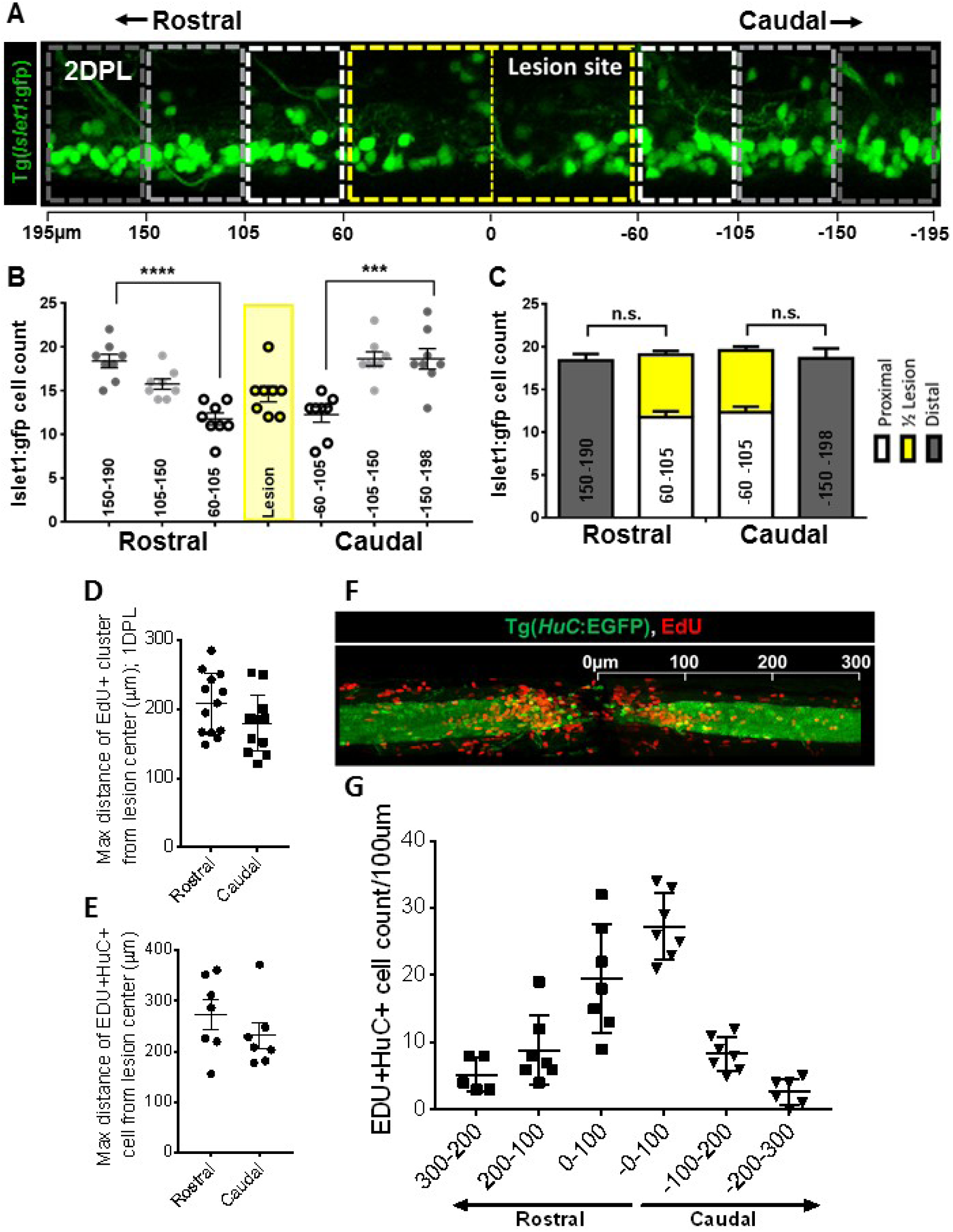
Neuronal precursor recruitment results in neurogenic response to replace cells lost both due to injury and as a result of migration. (A) 2DPL *Tg(Islet:GFP*) larvae with neighboring spinal segments outlined in dashed lines, lesion site outlined by yellow, these segments are used for subsequent analysis. Scale bar indicates the distance in µm from the lesion center (0), rostral segments are positive values while caudal values are negative. Each rostral and caudal segment is 45µm along the length of the spinal cord, while the lesion site encompasses a total of 120µm. (B) Total *Islet1*:GFP^+^ cell count within defined segments, the lesion site segment is outlined in yellow. For both rostral and caudal, the segment proximal to the lesion site had a significantly reduced cell count compared to the most distal segment. (C) A comparison of segment cell counts within both rostral and caudal stumps shows no significant difference between the distal segment *Islet1*:GFP^+^ cell count and the total cell count of the combined proximal segment and halved lesion site segment. (D) Mean maximal distance of EdU proliferation cluster from the lesion center at 1DPL was 208µm and 180µm for rostral and caudal spinal segments respectively. Clusters are defined as 3 or more cells within 10µm of each other. (E) Mean maximal distance of new born neuron indicated by EdU and HuC co-labelling at 6DPL was 274µm and 232µm for rostral and caudal respectively. (F) Representative maximal projection of 6DPL cumulative EdU pulse/chase in *HuC:EGFP* expressing larvae, scale bar of 100µm increments. (G) Quantification of the number of newborn neurons co-labelled with EdU and *HuC:EGFP* within each 100µm segment from the lesion centre reveals an inverse relationship between increasing distance from the lesion centre and number of newborn neurons.

To assess the range of neural stem and progenitor cell activation, the maximal distance of EdU^+^ cell labelling in the spinal cord was assessed at 1DPL, which was the time point when most widespread proliferation was detected (Figure S6 A and B). The maximal distance of proliferation was observed in the ependyma up to 208µm ± 12µm rostral and 180µm ± 11µm caudal from the lesion centre, respectively (Figure 7D). Interestingly, the mean distance of proliferation overlapped with the range of cell recruitment during migration suggesting that neurogenesis may be needed to replenish the depleted cells (Figure S6 C and D). To determine if the distribution and range of newborn neurons in response to injury correlated with the migratory and proliferative response, we performed cumulative EdU pulse chase experiments and assessed the co-localisation of EdU with the pan-neuronal marker HuC using immunohistochemistry at 6DPL. New born neurons indicated by EdU^+^/HuC^+^ co-labelled cells were detected up to 274µm ± 29µm and 232µm ± 25µm from the lesion centre rostrally and caudally (Figure 6E), overlapping with the range of cell migration and proliferation. When total numbers of EdU^+^/HuC^+^ cells were quantified within increasing segments of the lesion centre an inversely decreasing number of co-labelled cells was detected (Figure 7F and G). Taken together, the experiments demonstrate that the early neural precursor migration and depletion range correlates with the range and extent of injury induced cell proliferation and regenerative neurogenesis.

## Discussion

Tissue regeneration in animals takes place through two often cooperating modes: epimorphosis and morphallaxis (Morgan, 1901, Alvarado and Tsonis, 2006). Epimorphic regeneration is characterised by re-growth of lost tissue via cell proliferation and differentiation, while morphallactic regeneration is characterised by tissue remodelling and transformation of remaining tissue to replace the lost tissue. At present regeneration and restoration of tissue function in vertebrates is mainly thought of as stem and progenitor cell driven process following the epimorphic scheme (Tanaka and Ferretti, 2009, Poss, 2010). However, there are alternative less explored routes for tissue repair such as injury induced recruitment of dormant precursors or conversion of specific cell lineages into another.

We demonstrate that spinal cord regeneration in the zebrafish is driven by tissue remodelling in addition to activation of dormant stem and progenitor cells acting in concert during temporally distinct phases. Initial recovery is governed by tissue remodelling characterised by rapid cell migration of neural precursors from areas proximal to the injury, followed by an overlapping but later phase driven by cell proliferation and neurogenesis that restores lost tissue. Both processes are intimately linked on a cellular level, since the proximal region where depletion takes place, spatially overlaps with the extent of proliferative ERGs and subsequent regenerative neurogenesis. The two phases, neural precursor recruitment followed by compensatory regenerative neurogenesis, enable the zebrafish to rapidly regain motor function but also gradually replenish lost cells over time. Surprisingly, we find that that neural precursor recruitment and not cell proliferation driven neurogenesis contributes to the initial functional recovery. This is consistent with previous studies that reported very few newly produced neurons but substantial increase of cells with neural precursor characteristics such as *Islet1*:GFP or *Hb9*:GFP expression at lesion site during the first days and weeks following injury in larval and adult zebrafish respectively (Reimer et al., 2008, Briona and Dorsky, 2014, Ohnmacht et al., 2016). Furthermore, our findings are supported by a recent study in the adult zebrafish telencephalon which showed that a small population of neural precursors directly migrates to the lesion site and participates in repair without undergoing proliferation (Barbosa et al., 2015). Taken together, this suggests that recruitment of neural precursors may be a common mechanism enabling rapid neural repair leading to functional recovery in zebrafish.

We argue that the utilisation of existing neural precursors in CNS repair offers several benefits in addition to neurogenesis. The time-frame required to initiate and execute neurogenesis is too slow to acutely adapt or repair neural circuits at the cellular level. This is evident in the zebrafish CNS where it takes many days to weeks to replace neurons in the larval and adult zebrafish after injury (Kroehne et al., 2011, Kaslin et al., 2017, Reimer et al., 2008, Goldshmit et al., 2012, Hui et al., 2015). In the adult zebrafish spinal cord it takes many weeks to robustly detect newly produced ventral interneurons and motoneurons after lesioning demonstrating that neurogenesis driving regeneration is considerably slow (Reimer et al., 2008, Hui et al., 2010, Kuscha et al., 2012b). Slow neurogenesis and recovery is even more evident in mammalian models and in particular when human cells have been used to repair injury. For example, grafted human cells in rat spinal cord mature over many months and functional recovery is detected only after one year in line with the slow differentiation process (Lu et al., 2017). Using an endogenous reservoir of neural precursors, which are ‘ready to go’, provides efficient means to rapidly initiate tissue repair allowing time for the slower regenerative processes such as neurogenesis to commence. The recruited precursors may directly replace lost cell types but may also be important in re-establishing tissue continuity and permissive cues that promote neurogenesis and regrowth of axons. Consequently, it is important to further understand the role and mechanisms controlling such neural precursors since they could open up new strategies to facilitate neural repair in future therapies.

Neural precursors with immature characteristics have been shown in the CNS of multiple vertebrates including human and thought to represent cells that arrest or slow down their differentiation during neurogenesis (Kempermann, 2008, Marichal et al., 2009, Bonfanti, 2011, Kempermann, 2012). Such neural precursors have been proposed to act as a cellular reservoir to rapidly adapt neural circuits to cognitive challenges or injury (Kempermann, 2012). Intriguingly, a precursor population described as immature neurons in “standby mode” have been reported in the spinal cord of turtle and rat (Russo et al., 2004, Marichal et al., 2009) and other studies shown recruitment of precursors with neural characteristics after injury implying a possible role in repair (Mothe and Tator, 2005, Danilov et al., 2006, Shechter et al., 2007, Meletis et al., 2008). The standby cells in turtle and rat are distributed in the ependyma of the central canal, express early neuronal and plasticity markers such as HuC/D, PSA-NCAM, and exhibit electrophysiological characteristics of maturing neurons (Russo et al., 2004, Marichal et al., 2009). Similarly, we detect expression of HuC and PSA-NCAM in ependymal cells and cells recruited to the injury site during homeostasis and after injury. Intriguingly, delayed neurogenesis and neural precursors have recently been found in diverse brain regions of mammals (Luzzati et al., 2014, Piumatti et al., 2018, Rotheneichner et al., 2018). Furthermore, enigmatic cells with immature neuronal characteristics have frequently been reported after injury in diverse mammalian models (Lindvall and Kokaia, 2015). This suggests that maintaining cells with immature neuronal characteristics in the CNS is a conserved trait of vertebrates and contributes to repair. Taken together, our work demonstrates a role of precursor neurons recruitment in spinal cord regeneration and highlights a less explored route in neural repair.

## Material and Methods

### Animals

Fish were housed and bred within the Monash Fishcore Facility according to standard procedures. All experiments were approved by the Monash Animal Services Animal Ethics committee, Monash University. The following fish strains were used; *Tg(gfap:GFP*)^*mir2001*^, *Tg(HuC:EGFP*, HuC aka *elavl3*)^*knu3*^, *Tg(isl1:GFP*)^*rw0*^ and *Tg(−3.0mnx1:mTagBFP*)^*mq10*^. Clutch siblings were consistently used as controls and all embryos and larvae were maintained at 28ºC throughout development and experiments. Sex was randomly assigned to experimental groups.

### Larval spinal cord lesion assay

3dpf larvae were anaesthetized in 0.168mg/mL ethyl-m-aminobenzoate methanesulfonate (tricaine; Sigma) and embedded in 1% low melting point agarose on a 2% agar plated 90mm petri dish in a lateral position. An electrolytically sharpened 0.38mm tungsten wire needle (World Precision Instruments) was used to perform mechanical lesions at the level of the anal pore. Lesions were approximately one myotome, 100μm, in diameter and the notochord was left intact. Lesions were performed under an MVX10 Leica fluorescence microscope to ensure the complete removal of spinal cord tissue within the lesion site. Following the lesion larvae were gently freed from the agarose and placed into fresh RINGER solution (116mM NaCl, 2.9mM KCl, 1.8mM CaCl, 5mM HEPES in MilliQ to pH7) containing 100μg/mL penicillin/streptomycin (Sigma). Larvae displaying damage to the notochord were removed from the experiment.

### Adult spinal cord lesion assay

Adult fish were lesioned as previously described (Goldshmit et al., 2012), with the exception that the wound was not sealed. Following injury fish were monitored daily and deaths were immediately removed from tanks.

### Quantification of percent regeneration

Two alternative methods of percent regeneration quantification were applied based on the transgenic expression used for analysis. *Tg(gfap:GFP*) is expressed by stem and progenitor cells, however due to the persistent perdurance of GFP in differentiating cells this label is broadly detected within the larval spinal cord. In addition, a high exposure was used to *acquire Tg(gfap:GFP*) images in order to detect subtle processes at the lesion site. Similarly, as an early marker of neuronal fate, *Tg(HuC:EGFP*) is broadly expressed throughout the spinal cord. Due to this broad expression the identification of single cells within *Tg(gfap:GFP*) and *Tg(HuC:EGFP*) acquired images was not possible. Therefore a maximal intensity approach to percent regeneration quantification was adopted (Figure S1, A-G). Using Fiji/Image J software stacks were compressed into maximal projection images and converted into binary format. In a duplicated window the lesion site was filled in to make a pseudo-continuous column from which a mask was created. The original binary image was then subtracted from the inverted format of the mask to give an impression of the size of the lesion site, from which an area measurement could be taken. This area of the lesion site then subtracted from the original lesion site size to give a quantification of re-growth. The percent regeneration was then calculated by dividing the re-growth area from the initial lesion size area, and multiplied by 100. Due to the distinct cellular localisation of the *Tg(Islet1:GFP*) reporter a cell counting approach was adopted for quantifying percent regeneration (Figure S1, H-M). The Imaris Cell Counter plugin was used to quantify cell numbers needed for this approach. Cells were counted within a region of interest defined by encompassing the entire z axis, all cells along the dorsal/ventral axis and all cells with 140μm along the rostral/caudal axis. The thresholding tool was used to ensure the spots registered matched the cells observed within the region of interest. The number of cells within the lesion site was recorded and the same region of interest and thresholding settings were used to calculate the cell number within the intact region rostral of the lesion site, within the same larvae. The rostral cell count provided an internal control to compare the number of cells in the lesion site to. Percent regeneration was calculated by dividing the total cell number from the lesion volume by the total cell number in the rostral volume and multiplying by 100.

### Locomotor function assessment

To evoke an escape response larvae were touched with a plastic micropipette on the tip of the head. The quality of locomotor function was measured by recording the maximal angle created by larvae during the initial escape reflex, known at the C-bend angle (Eaton et al., 1984, Kimmel et al., 1974, Fero et al., 2011). All C-bend angle recordings were acquired with a high speed camera capturing 1,000 frames per second using Stream Pix 5 recording software. Recordings were then analysed using ViewPoint software to measure the maximal C-bend angle. This was identified as the maximal angle produced between the tip of the tail relative to the central direction of the head.

### Muscle force recordings

Larvae were immobilised by incubating on ice for 5min, then euthanized and mounted horizontally using two aluminium clips onto an in vitro apparatus (model 1500A, force transducer 403A, Aurora Scientific, Ontario, Canada). One clip was placed immediately caudal to the lesion on the anterior side, and the other clip was wrapped at the end of the tail (Figure 2 D), therefore the force measured upon stimulation was from the trunk muscles in between the two clips. The preparations were held in a MOPS buffered physiological solution at 22°C and stimulated via two platinum electrodes (Li et al., 2013). The optimal length for active contraction was first determined, then muscles were kept at this length and stimulated using varied pulse durations (0.1–5 ms) to examine the excitability of the neuromuscular activation (Abramsson et al., 2013). For analysis, the active force measured at each pulse duration was normalised to the active force at 5ms as the maximal.

### GCaMP imaging and analysis

Zebrafish 4-6 dpf larvae were immobilized right side up in 2% low melting point agarose (Sigma-Aldrich) on microscope slides. The agarose surrounding the fish was cut vertically, parallel to the fish, with a scalpel, and removed. The embedded fish was transferred to custom made, 3D printed chamber. The chamber was filled with E3 media. Larvae were then transferred to the imaging room and allowed to acclimate for 15 min prior to imaging on a custom built diffusive lightsheet microscope (Taylor et al., 2018). Timelapse stacks were acquired at 500ms intervals for a period of 10 minutes. On average larvae had 3-5 spontaneous motor contractions within this time period. Motion artifacts caused by slow drift of the image or by spontaneous movements by the larva were corrected in Fiji using “moco” (Dubbs et al., 2016) followed by a rigid body transformation in StackReg (Thevenaz et al., 1998). ROIs were drawn by hand in Fiji to select the ventral part of the spinal cord, and the mean fluorescence of the ROI was plotted over time. The raw fluorescence time series were further analyzed in Matlab. ΔF/F was calculated as previously described (Jia et al., 2011), and the local maxima function from Matlab was used to identify peaks of activity rostral of the lesion site with a threshold of 3. The maximum intensity of the rostral activity spike was used to normalize the activity, during the same window of time, in the caudal part of the lesion site.

### 5-ethynyl-2’-deoxyuridine (EdU) labelling and detection

Larval labelling consisted of placing fish in 50μg/mL EdU and 10% DMSO in RINGER solution, with the duration of incubation dependant on experimental approach. To provide a snapshot of the amount proliferation occurring at each consecutive day after injury fish were briefly pulsed in EdU solution on ice for 30 minutes, followed by 30 minutes at 28ºC. Larvae that were 5dpf and older were removed from ice treatment at first sign of reduced circulation. Following the pulse larvae were immediately rinsed with RINGER solution to remove all DMSO and left to incubate under normal conditions for a further 2 hours. To identify proliferation leasing to neurogenesis fish were either incubated for up to 6DPL in normal conditions or underwent additional pulses in EdU solution as previously described. At the chase end point larvae were euthanized, the torso rostral to the yolk-budge was surgically removed and tails were fixed in 4% paraformaldehyde (PFA) in Phosphate Buffer (PB, Monosodium phosphate, Disodium phosphate, pH 7.4), overnight at 4 degrees. Following permeablisation in PBSTx (0.3% Triton 100 X in PBS) EdU labelling was detected by incubating tails in fresh EdU Click-iT reaction solution (Roche) for 2 hours in the dark at room temperature. Following three 30 minute washes in PBSTx tails were either mounted for imaging in 70% glycerol or subsequently processed for immunohistochemistry.

Adult labelling consisted of intraperitoneal injections of 10mM EdU as previously described (Lindsey et al., 2018). Fish were injected at 3, 5 and 7 days post injury and chased until 18 days post injury at which point fish were euthanized and the torso was dissected for fixation in 4% PFA in PB. The tissue was then permeabilised in EDTA solution (200mg/mL sucrose, 0.5M EDTA in 1x PBS) and mounted in embedding solution (200mg/mL sucrose, 50mg/mL fish gelatine in 1xPBS). For cryosectioning. EdU labelling was detected by incubating sections in fresh EdU Click-iT reaction solution (Roche) for 30 minutes in the dark at room temperature and were then processed for further IHC.

### Whole-mount immunohistochemistry

Larval tails were previously fixed and extensive washed in PBSTx prior to IHC. All incubations were performed at 4ºC for between 1 to 4 days. Tails were blocked in 1% DMSO, 2% normal goat serum (NSG; Sigma-Aldrich) in PBSTx. Primary antibody were incubated with 2% NGS. Following extensive washing in PBSTx tails were incubated in secondary antibody with 10% NSG, washed and then mounted for imaging in 70% glycerol.

### Drug treatments

To block cell proliferation the cyclin-dependant kinase (CDK) inhibitors Olomoucine and Ryuvidine (Sigma) where applied, blocking CDK2 and CDK4 activity respectively. Embryos were treated at 1 day post lesion with 20μM Olomoucine or 1μM Ryuvidine containing 0.5% DMSO in RINGER solution and incubated for 24 hours. To block cell migration larvae were incubated with 50ng/mL Nocodazole (Sigma) with 0.5% DMSO in RINGER solution at 1 day post lesion and incubated for 24 hours. Larvae we incubated at 28ºC, following incubation larvae were imaged and assessed for percent regeneration or contractile force production. When assessing GCaMP6s responsiveness larvae were incubated in Nocodozole from 0 to 3 days post lesion. To selectively block tubulin formation at the lesion site injured larvae were incubated in 50µM photostatin (donated by Dr. Oliver Thorn-Seshold) and subjected to a 405 laser during the first 24 hours following injury. Laser intensity was kept low, 0.5%, and 15µm steps were applied through the width of the cord at 15 minute intervals throughout the timelapse. Injured control larvae were mounted within the same dish, containing the photostatin solution, but were not subjected to laser treatment.

### Image acquisition and processing

All confocal fluorescent images were acquired on either a TCS SP8 or TCS SP5 confocal (Leica Microsystems) with a Leica 20x/1.00NA HCX APO objective. Stacks were acquired with 1μm step size and encompassed the entire transverse aspect of the spinal cord. Files were viewed in Imaris (Bitplane) and final images were saved as tiffs and exported to Powerpoint (2013) and cropped for final presentation.

### Spinal cord cell tracking and registration strategy

Quantitative analyses of neuronal cell migration during spinal cord recovery were performed using custom scripts and an extension for Bitplane Imaris programmed in Python using the numpy (Walt et al., 2011), scipy and scikit-image (Walt et al., 2014) libraries. The source code for these are available at https://gitlab.erc.monash.edu.au/skeith/scr and https://github.com/keithschulze/xtscr.

To assess the migration of neuronal cells along the length axis of the spinal cord, we modelled the spinal cord as a cylinder where the position of each cell is defined by coordinates along the cylindrical axes: length (l), radius (r) and angle (θ, Figure S4A). To determine the cylindrical coordinates of each cell, we first needed to determine the central axis of the spinal cord, which corresponds to the length axis of the cylinder. During imaging, fish were orientated such that the spinal cord was approximately aligned along the Y axis of the image. Sub-volumes spanning the entire XZ dimensions and 10 pixels in the Y dimensions were extracted from the caudal and rostral ends of the Y axis. These regions were then max intensity projected in Y, such that 2 max intensity XZ slices from the extremities of the Y axis were created. A 10-pixel Gaussian blur filter was applied to smooth the images. The spinal cord region in both images was then segmented by applying an Otsu threshold (Otsu, 1979) and selecting the largest labelled region in the image. Bounding boxes for these labelled regions were used to subset the smoothed max intensity projected images to extract just the areas containing fluorescence intensities from the spinal cord. These subsetted regions roughly correspond to the max intensity projection of the boxes highlighted in yellow in Figure S4B. These subsetted regions were then fitted with the following 2D Gaussian function:

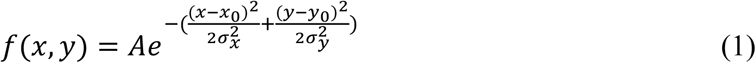

where A is the amplitude, *xx*0,*yy*0 are the coordinates of the centre of the peak and *σσxx,σσyy* are the standard deviation (i.e., width) of the tail in the respective dimensions. Note that the tails in *xx* and *yy* dimensions were allowed to vary independently to allow for a non-symmetrical Gaussian fitting. The coordinates for the centre of each Gaussian fitting (i.e., *xx*0,0) were designated as the start and end coordinates of a line running down the central axis of the spinal cord. An example of fitting a Gaussian to the max projected intensity images at the caudal and rostral ends of the Y-axis is shown in Figure S5C (left). A nice property of this approach is that the determination of the central axis was quite resistant to part of the spinal cord drifting out of the field of view (Figure S5C, right).

To register the spinal cords from different datasets to a common reference frame, we manually specified coordinates for the centre of the lesion site and the centre of the adjacent notochord by placing Bitplane Imaris “Measurement Points” at these locations. The Measurement Point (MP) placed at that centre of the lesion site was then projected onto the central axis and used to define the origin of the length (l) axis, where negative values were caudal to the lesion centre and positive values were rostral to the lesion centre. All cell coordinates were then transformed such that θ = 0° for the MP placed in the notochord i.e., spinal cord datasets were rotated around the length axis until the centre of the notochords were aligned.

To analyse movements along the length axis of the spinal cord, the length coordinate at each timepoint of a given cell track was extracted and the time series smoothed using a Savitsky-Golay filter (Savitzky and Golay, 1964) with a window size of 5 and polynomial order of 2. Smoothed tracking data was then exported and plotted using R (Ihaka and Gentleman, 1996) and the ggplot2 package. Normal distribution and sample size. We tested if our quantifications of regeneration (motorneuron or glia) follow normal distribution using Q-Q plots & Shapiro-Wilk’s test and determined our data to be normally distributed (α<0.05, p<0.05). Using power calculations for 80% power at 5% significance level and S.D. values measured from experiments yield that 4-6 animals are required. Consequently we have adopted cohorts of 5-10 fish/experiment.

### Quantification and statistical analysis

All experiments were performed with a minimum of 3 biological replicates and repeated independently at least twice; exact numbers are shown in figures. Statistical analysis was performed with Prizm (GraphPad) using student’s unpaired two-tailed t test when comparing two conditions. ANOVA with Tukey’s post hoc analysis was used when comparing multiple samples with each other.

## ACKNOWLEDGEMENTS

Authors thank Monash Micro Imaging (MMI) facility, Monash University, for technical imaging support and Dr. Thorn-Seshold for generously donating the photostatin used in this manuscript. We would also like to thank AquaCore, Monash University, for microscope access and support with fish husbandry. This project was funded by NHMRC project grants GNT1068411, GNT1145048 and GNT 1138870 (JK); HC was supported by fellowships from CNPq (202130/2015-0) and Sao Paulo Research Foundation (FAPESP – 2017/06022-7). The Australian Regenerative Medicine Institute (ARMI) is supported by operational Infrastructure from the Victorian Government.

## AUTHORS CONTRIBUTIONS

J.K. and C.V. conceived experiments. C.V. conducted all experiments for the study and H.C., M.L., M.K. and G.V assisted with tissue processing, force recordings, immunohistochemistry, and/or confocal imaging. C.V. analysed all data sets with assistance from G.V, M.L. and K.S. J.K and C.V drafted figures and manuscript with input from H.C and M.A All authors reviewed and edited the manuscript for final submission and revisions.

---

**Figure S1.**
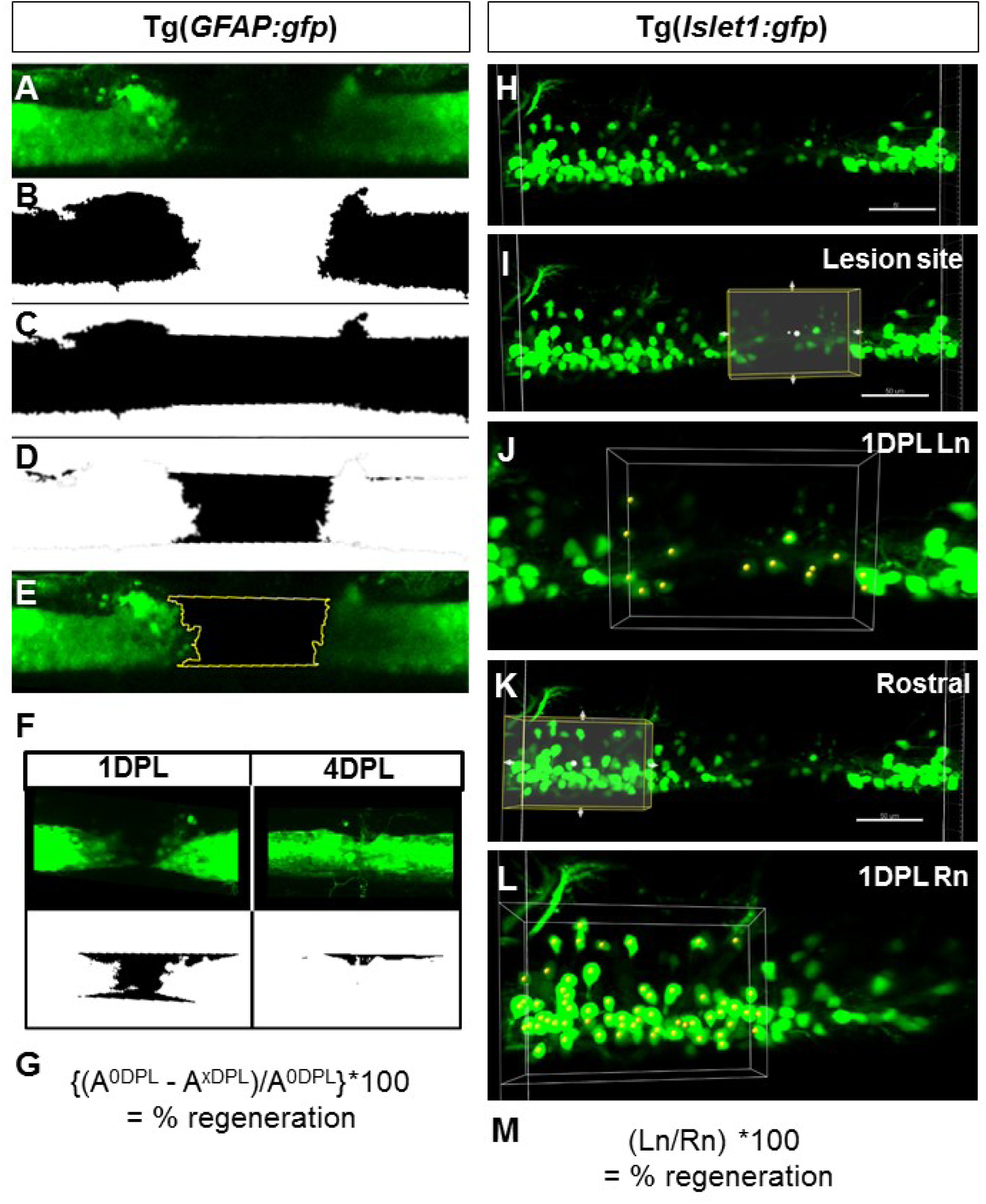
Related to Figure 1. Quantification of percent regeneration. Image of pan-neuronal reporter *Tg(HuC:EGFP*) at 0DPL used as example of quantifying initial lesion size. (A) Using Fiji/Image J software confocal stacks are converted into a maximum projection image. (B) Image is converted to binary. (C) On a duplicated image the polygon selection tool is used to fill the lesion gap. (D) The Image Calculator tool is used to subtract the original binary image (B) from pseudo-filled binary image (C), leaving an area replicating the lesion size which is measured using the selection tool. This area value is the A0DPL measurement used to quantify percent regeneration at later timepoints. (E) The outlined selection of the lesioned area conforms to the lesion site in the original maximum projection. (F) Maximum projections of *Tg(gfap:GFP*) at 1 and 4DPL along with the corresponding binary lesion size areas as measured using steps A-D. A distinct reduction in the lesion area is observed over time. (G) Percent regeneration is calculated by subtracting the lesion area of a given time point after injury (AxDPL) from the initial lesion area (A0DPL), then dividing this total from the initial lesion area (A0DPL) and multiplying by 100. (H) Image of motor neuron reporter *Tg(Islet1:GFP*) at 1DPL is used as an example. The Cell Counter plugin within Imaris (V8.4) software is used for counting cells. (I) A region of interest (ROI) is defined by encompassing the entire z axis, all cells along the dorsal/ventral axis and all cells with 140µm along the rostral/caudal axis. The ROI volume (yellow box) is placed over the lesion site and thresholding for Quality is adjusted to overlap with *Islet1*:GFP expressing soma, indicated by yellow spheres (J). The number of identified cells is recorded as Ln. (K) Using the same Quality thresholding value the ROI volume is placed rostral to the lesion site to gain the number of *Islet1*:GFP expressing soma within intact regions of the spinal cord, indicated by yellow spheres (L). The number of identified cells is recorded as Rn. (M) The percent regeneration is then calculated by dividing the total cell number from the lesion volume (Ln) by the total cell number in the rostral volume (Rn) and multiplying by 100.

**Figure S2.**
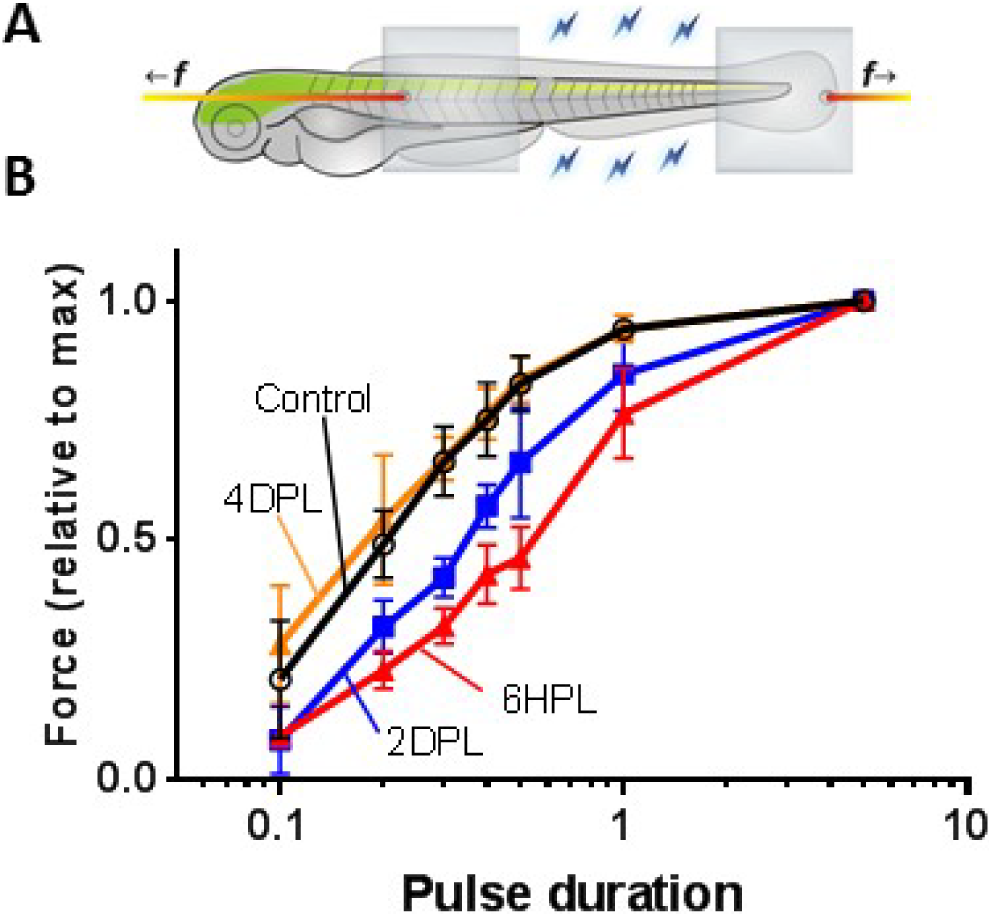
Related to Figure 2. (A) Schematic of muscle force recording preparation with aluminum clips indicated by grey boxes, between which force measurements are recorded as indicated by blue bolts. Orange lines with f indicate the mechanical pull exerted on larvae to maintain optimal muscle length. (B) Relative force generation by the muscles caudal of the lesion site under varied electrical stimulations (pulse duration range of 0.1-5ms) in uninjured control, 6HPL, 2DPL and 4DPL groups. Relative force production curve was reduced for both 6HPL and 2DPL compared to 4DPL and control groups. N=4 larvae per group.

**Figure S3.**
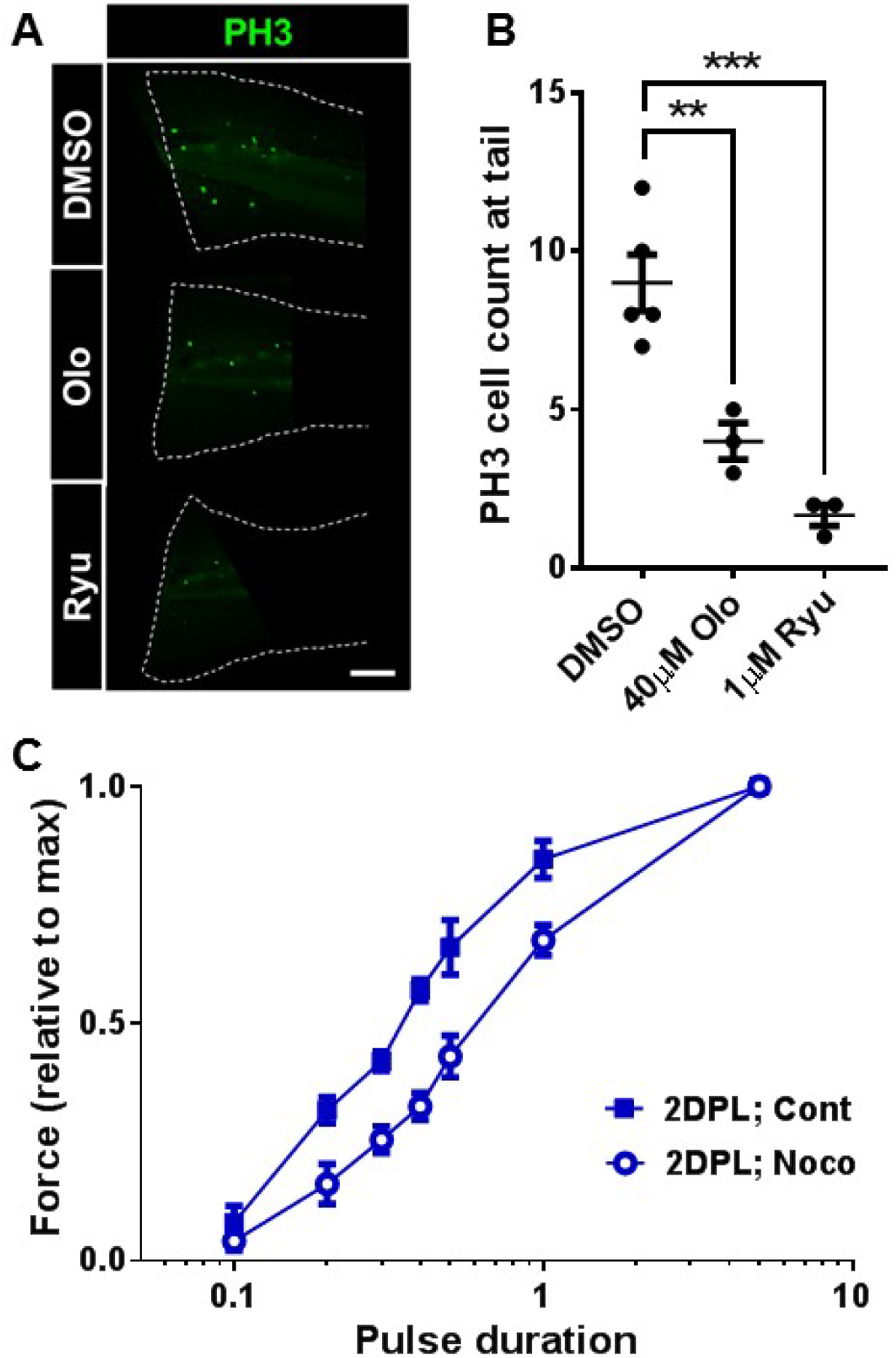
Related to Figure 5. Tail snip was performed on 3dpf larvae to induce a proliferative response, which was followed by immunostaining for the proliferative marker PH3. (A) Representative widefield fluorescent images of injured tails at 1DPL in control and treatment conditions, scale bar = 100µm. (B) Quantification of the number of PH3 positive cells within the portion of the tail caudal to the end of the notochord. Compared to control the number of PH3 cells was significantly reduced with both 40µM Olomoucine and 1µM Ryuvadine treatments. (C) Force production curve in control and 50ng/mL Nocodozole treated larvae at 2DPL. The overall generation force was significantly reduced with Nocodazole treatment. N=3-5 larvae, error margin = SEM (standard error of the mean), p= **<0.01, ***<0.001. Statistical analysis for B was measured using students t-test.

**Figure S4.**
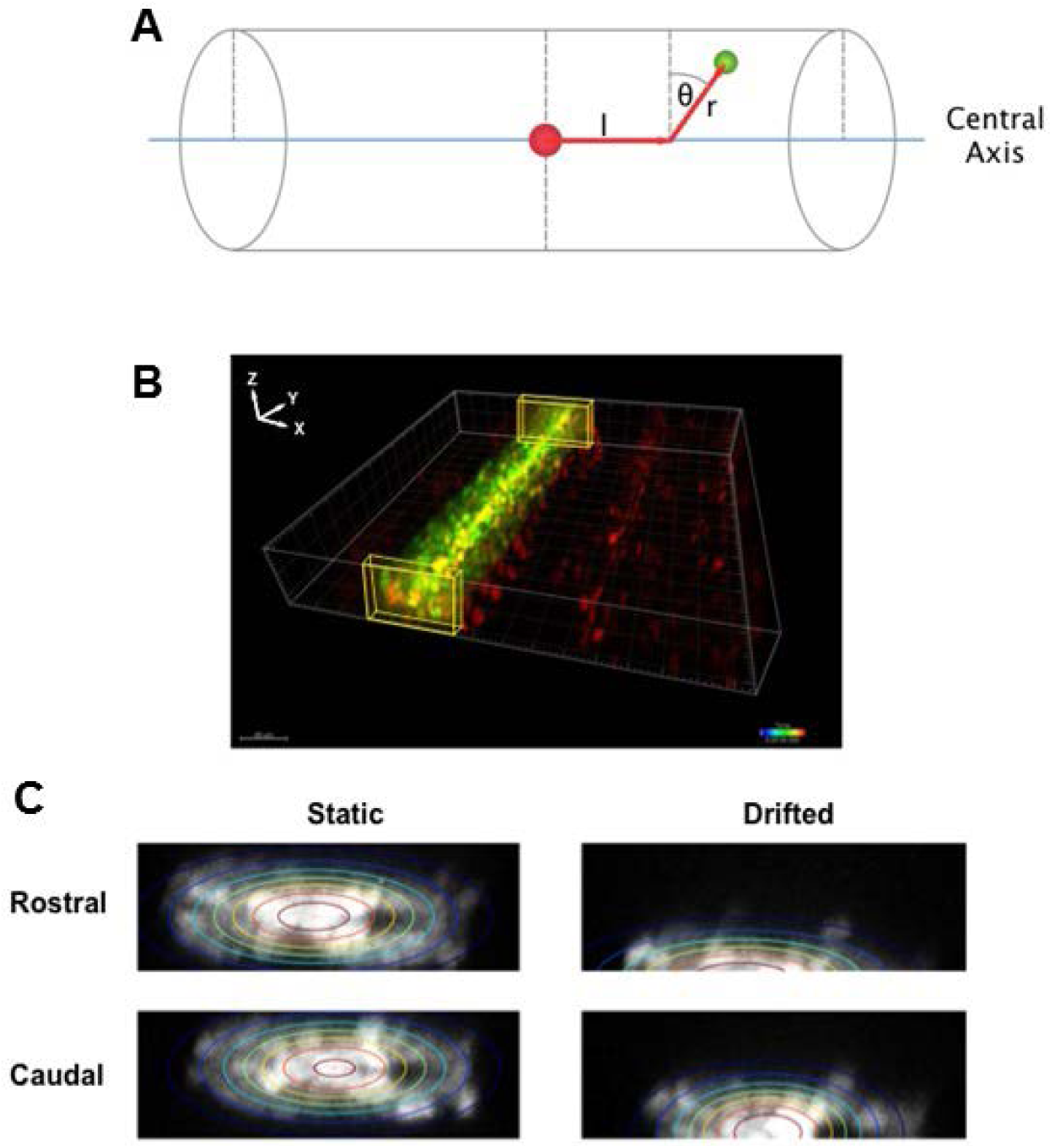
Related to Figure 5. Strategy for fitting the central axis of cylindrical coordinate system for the spinal cord. (A) Schematic of cylindrical coordinate system. (B) Sub-volumes (yellow boxes) extracted to produce maximum intensity projections for the rostral and caudal limits of the spinal cord in the Y-Axis of the image. Two-dimensional Gaussian functions fitted to smoothed maximum projected image intensity data from the rostral and caudal limits of the spinal cord in the Y-axis. The centre of the fitted gaussians (rostral and caudal) is used to approximate the coordinates of the limits of a line running down the centre of the spinal cord. (C) Schematic of cylindrical coordinate mapping of cells in the spinal cord.

**Figure S5.**
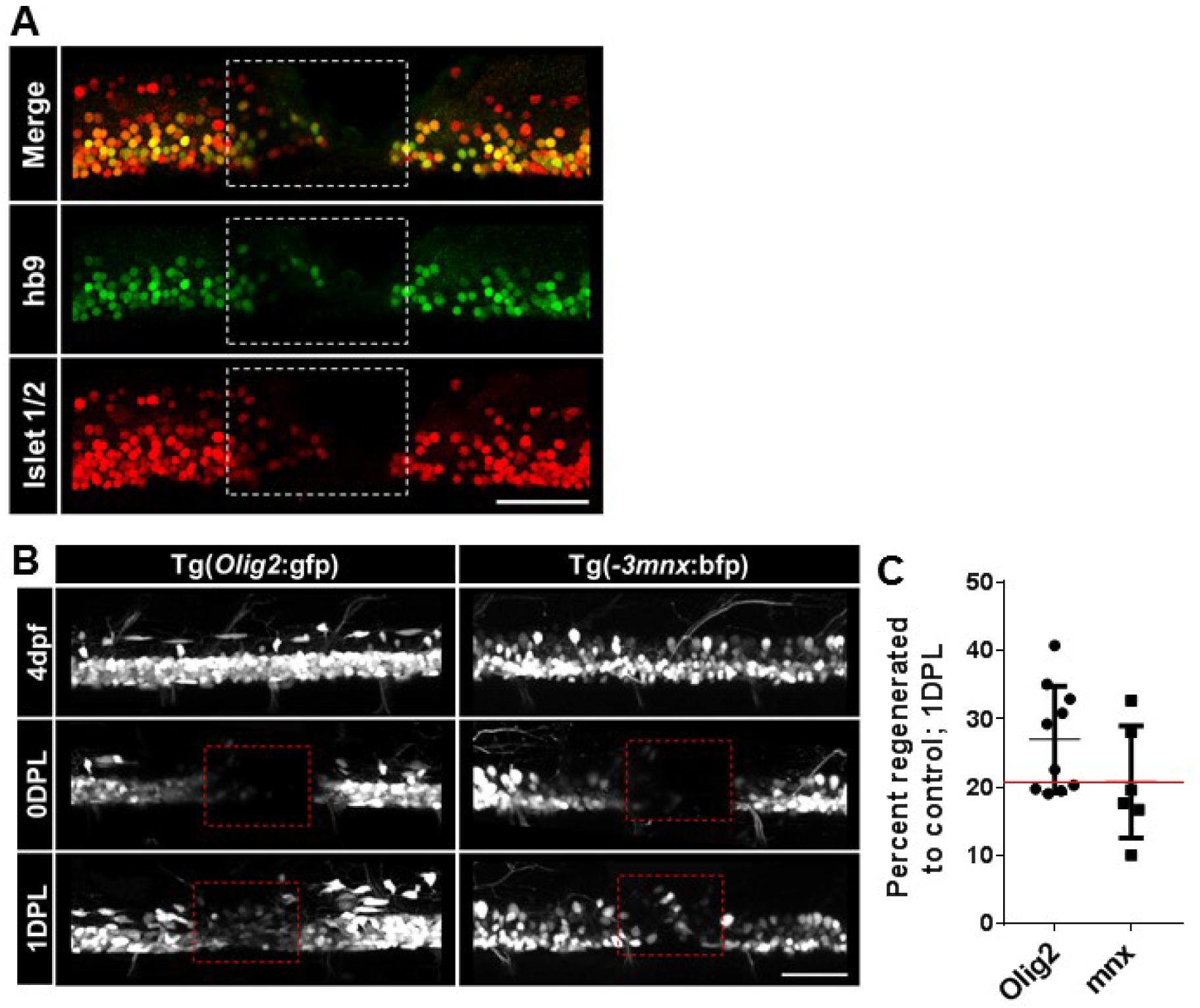
Related to Figure 6. (A) Immunohistochemistry in 1DPL larvae against Hb9 and Islet1/2 proteins. Merge panel shows many cells within the lesion site express both markers. (B) Maximum projection of confocal images within the *olig2*:EGFP and *mnx1*:BFP transgenic lines at 4dpf, 0DPL and 1DPL. 0DPL images show a lack of cells within the lesion site, indicated by red lined box. At 1DPL both *olig2*:EGFP^+^ progenitors and *mnx1*:BFP^+^ neurons are located within the lesion site. (C) Quantification of cellular regeneration seen in (B). Red line indicated mean percent regeneration of *Islet1*:GFP^+^ cells at 1DPL, as quantified in Figure 1P. Scale bar = 100µm. Dorsal up and rostral to left in all panels.

**Figure S6.**
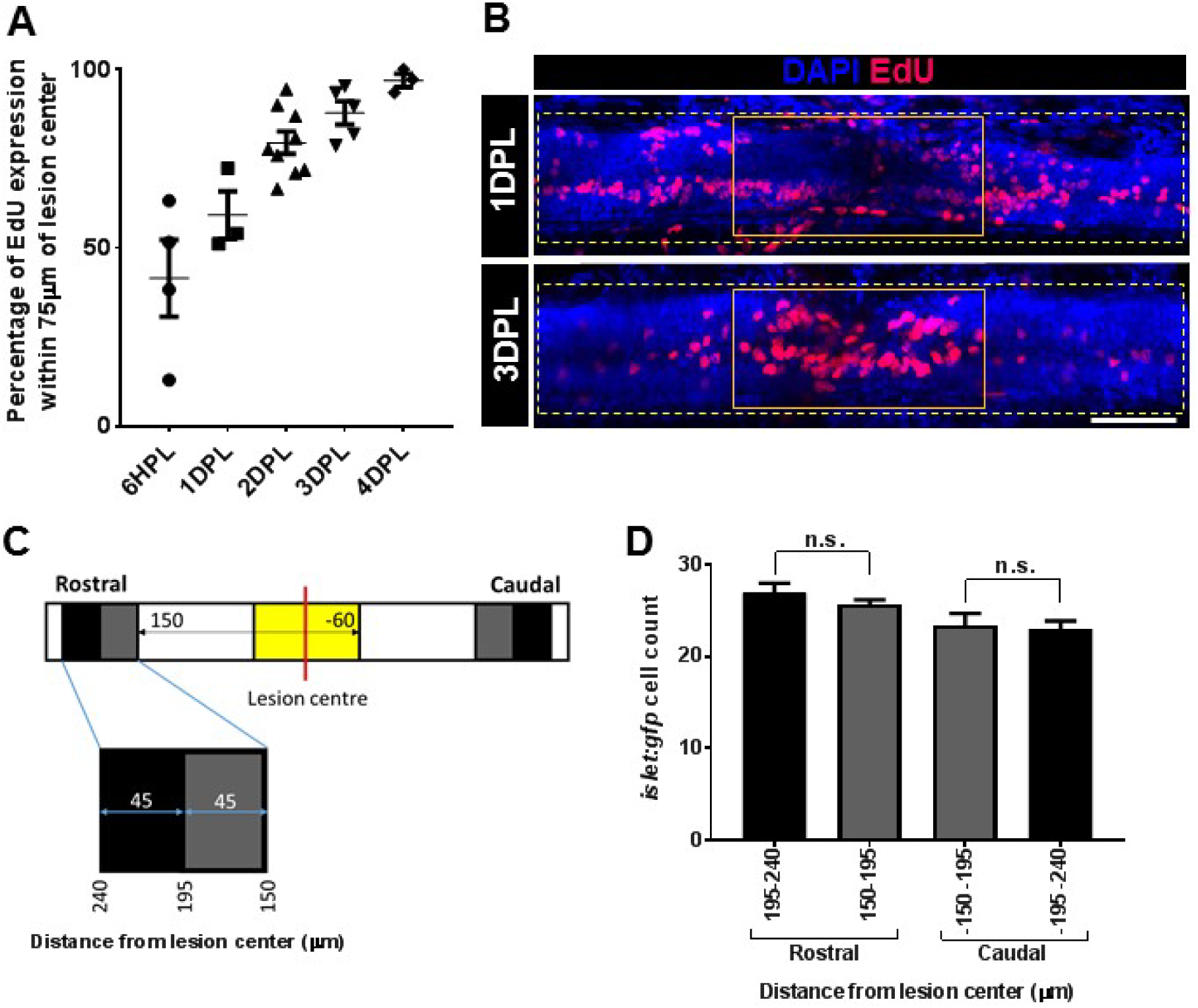
Related to Figure 7. Extent of proliferation and migration is restricted along the A-P axis. (A) Quantification of percent of EdU cells within the spinal cord being localised within 75µm rostral and caudal of the lesion centre compared to the entire spinal cord captured during confocal imaging. Over time the percentage of EdU cells with 150µm of the lesion centre increases, indicating that proliferation becomes restricted to the local lesion site. (B) Representative maximal projections of confocal images taken at 1DPL and 3DPL. EdU cells are largely within the solid orange box outlining 150µm surrounding the lesion centre at 3DPL compared to cell being more distributed within the larger yellow dashed box at 1DPL. (C) Distal regions for analysis were measured within 45µm neighbouring segments that begun at a distance of 150µm from the lesion centre. (D) No significant difference between the total *Islet1*:GFP^+^ cell number is detected within neighbouring distal segments, both rostral and caudal of the lesion centre. n.s. = not significant, n= 3-12. Scale bar = 50µm.

## Supplementary movies

**Movie S1**. **Rapid glial bridging.** Time lapse of initial glial regeneration starting at 2HPL and ending at 24HPL. *Tg(gfap:GFP*) displayed in grey scale. Stacks were acquired every 18.5 min using a 20X 1.0NA objective on a Leica SP5 confocal microscope. Rostral to the left and dorsal is up.

**Movie S2. Return of touch evoke response after injury.** Video recordings of the touch evoked escape response in injured and control larvae. Larvae are stimulated by touching the head with a tungsten needle. Larvae produce their maximal C-bend angle at 0.45 seconds after the response is elicited. This is followed by an acceleration away from the stimulus in both sham and 2DPL larvae. After 3 seconds both sham and 2DPL larvae have nearly swum out of the field of view. Larvae that are 6HPL lack an acceleration burst following the initial C-bend and remain close to the stimulus after 3 seconds. Video recording playback is 0.45 seconds per second.

**Movie S3. *Islet1*:GFP cell migration following injury.** Time lapse of initial *Tg(Islet1:GFP*) regeneration starting at 2HPL and ending at 24HPL. Stacks were acquired every 7.5 min using a 20X 1.0NA objective on a Leica SP5 confocal microscope. Rostral to the left and dorsal is up.

